# Evidence of active phage replication during vegetable fermentation and implications for the dynamics of bacterial communities

**DOI:** 10.64898/2026.10.02.756237

**Authors:** Julia Cantuti Gendre, Margareth Renault, Anne-Cécile Leforestier, Gwendoline Marquet, Françoise Irlinger, Nathan Fouchard, Sophie Landaud, Claire Le Marrec, Stéphane Chaillou, Eric Dugat-Bony

## Abstract

Spontaneous vegetable fermentation is driven by the indigenous microbiota associated with the raw ingredients. The process often involves an initial heterofermentative phase followed by a predominantly homofermentative phase, both ensured by lactic acid bacteria (LAB). While microbial dynamics during fermentation are well characterized, phage communities remain poorly understood, despite evidence linking phage activity to LAB succession. We monitored viral and bacterial succession during sauerkraut and carrot fermentations using a metagenomics approach to characterize community dynamics. To further assess the impact of phage PLE001 on bacterial community dynamics, we developed a simplified carrot fermentation model comprising representative strains of Rahnella aquatilis, Leuconostoc mesenteroides and Lactiplantibacillus plantarum. Two consortia were assembled: one containing the PLE001-susceptible strain L. mesenteroides CIM3004, and the other containing also its phage-resistant derivative, CIM3004 BIM. We observed major shifts in the viral community during the first week of fermentation, coinciding with changes in bacterial structure during hetero- and homofermentative phases transition. In the simplified model, phage PLE001 reduced the population of the host during the heterofermentative phase but did not affect overall fermentation dynamics nor metabolite production. These results suggest that bacteriophages contribute to bacterial succession during fermentation, while communities remain resilient to phage infection, preserving fermentation stability.

## 2. Introduction

Bacteriophages, or phages, are bacterial viruses that play a key role in shaping, regulating and maintaining the stability of microbial community structure and function (Brown et al., 2022). Bacteriophage ecology is a growing research field aiming to elucidate how these viral particles interact with other microbes, macro-organisms, and abiotic factors in the environment (Burmeister et al., 2019). Their importance as regulators of microbial communities was first highlighted in the ocean, where marine viruses lyse up to 20% of the microbial biomass daily (Suttle, 2007). Since then, the ecological roles of phages have been explored across diverse environments, ranging from natural ecosystems, such as soils (Braga et al., 2020), to anthropogenic environments such as fermented foods (Paillet and Dugat-Bony, 2021).

Fermented foods, including dairy products, meat, fish, seafood, vegetables, legumes, pulses, cereals, are produced through the metabolic activity of beneficial microorganisms, such as bacteria, yeasts, and molds (Tamang et al., 2020). Food fermentation is a highly dynamic process characterized by the sequential development of distinct microbial populations, ultimately resulting in unique sensory and nutritional properties. The presence of phages is well-documented in the dairy sector, where they can significantly affect product quality (de Melo et al., 2018; Ortiz Charneco et al., 2023; Samson and Moineau, 2013). However, numerous phages have also been isolated from plant-based fermentations such as sauerkraut, kimchi, and cucumbers (Barrangou et al., 2002; Yoon et al., 2002; Kleppen et al., 2012; Lu et al., 2012). These culture-dependent studies confirmed the presence of phages infecting the most dominant lactic acid bacteria (LAB) driving the fermentation, including Leuconostoc, Lactiplantibacillus, Levilactobacillus, Weissella and Pediococcus species. This body of knowledge has recently been expanded through the application of viral metagenomics, which enables the detection of bacteriophages without prior culture of bacterial isolates. This approach revealed a wide and unexpected diversity of viruses present in fermented vegetables, primarily comprising LAB and non-LAB bacteriophages (Cantuti Gendre et al., 2025; Jung et al., 2011; Park et al., 2011).

Confirming the ecological role of phages as key regulators in the assembly and functioning of microbial communities in fermented vegetables requires direct or indirect evidence of active phage replication during the fermentation process. Indirect evidence can be inferred from longitudinal studies describing the temporal dynamics of natural phage and microbial populations using metagenomic approaches. Lu et al. already reported, using isolation-based techniques, the succession of at least two phage-host systems during the fermentation of commercial sauerkraut (Lu et al., 2003) and cucumbers (Lu et al., 2012) in the USA, corresponding to a shift from heterofermentative (Leuconostoc, Weissella) to homofermentative (Lactiplantibacillus, Levilactobacillus) LAB populations. This initial view was later expanded by a viral metagenomics study of kimchi, which supported the temporal succession of distinct phage populations during fermentation and their association with compositional changes of the bacterial community (Jung et al., 2022). Results indicated that phages could reduce the population density of superior competitors while simultaneously increasing the population density of weaker competitors, as predicted by the theoretical “kill the winner” hypothesis (Thingstad, 2000). However, the temporal dynamics of phage populations in vegetable fermentations, and their associations with bacterial successions remain largely unexplored, particularly in spontaneously fermented products consumed in Europe.

Providing direct evidence of phage replication during spontaneous vegetable fermentation is challenging due to the complexity of natural microbial communities. Synthetic microbial ecology, defined as the use of simplified microbial communities assembled under controlled conditions, is an emerging field at the intersection of synthetic biology and microbial ecology, offering a powerful framework to overcome this limitation (De Roy et al., 2014; Großkopf and Soyer, 2014; Kong et al., 2018). Model microbial communities are commonly used to study microbial interactions in fermented foods (Mounier et al., 2008; Niccum et al., 2020; Pourcelot et al., 2025). However, only a few studies have explored the role of bacteriophages as modulators of food microbial communities using synthetic ecosystems (Spencer-Drakes et al., 2024). Notably, Erkus et al. examined the impact of phages on the dynamic of bacterial populations in an undefined starter culture used for cheese production in a laboratory-scale setup mimicking the backslopping procedure (Erkus et al., 2013). This research suggested that phage parasitism maintained diversity by suppressing the most abundant strains, thereby stabilizing the overall community’s functionality. Recently, Spus et al. pursued these investigations using a synthetic community representative of the same undefined starter culture, propagated in milk for over 500 generations and challenged with a cocktail of virulent phages (Spus et al., 2023). Finally, Ledormand et al. provided further insights into the role of phages during LAB-driven malolactic fermentation in cider (Ledormand et al., 2022). This work reinforced the idea that phages play an essential role in microbial communities, as they can modulate both microbial populations and their associated metabolites. To the best of our knowledge, such an approach has not yet been implemented to study the impact of phages on the microbial ecology of fermented vegetables.

In this study, we aim to provide both indirect and direct evidence of active phage replication during vegetable fermentation. We first investigated the temporal dynamics of phage and bacterial populations during lab-scale spontaneous fermentation of sauerkraut and carrots, using shotgun and viral metagenomic approaches. By demonstrating the concomitant variations of phages and their bacterial host, our results provide indirect evidence of phage-host interactions during fermentation. We further explored this phenomenon by establishing a synthetic ecology experiment to examine the impact of a deliberately introduced virulent phage on a simplified microbial community that mimics that of fermented carrots. Classical fermentation indicators, including pH, titratable acidity, lactic acid concentration, phage and bacterial counts, were monitored at three time points during fermentation.

## 3. Materials & methods

### 3.1. Laboratory-scale spontaneous vegetable fermentation

#### 3.1.1. Experimental setup

Fresh organic carrots and cabbages were collected from a farm located in the Grand-Est region in France. Vegetables were washed, grated, vacuum-packed separately, then stored at 4°C until use. A single batch of laboratory-scale fermentation was prepared for each type of vegetable separately (Supplementary Figure S1). Briefly, 3.2 kg of grated vegetables were mixed with 48 g of sterile sea salt (1.5% w/w) in a sterile beaker placed under a biological safety cabinet, and homogenized manually using sterile gloves. The mixture was incubated for 1 h at room temperature before being distributed into 23 jars, each containing 135 g of product. To ensure an even distribution of vegetable and exudate among all the jars, the exudate was first collected by manually pressing the mixture with sterile gloves and gauze, then weighed. The remaining vegetable was also weighed. Both the vegetable material and the exudate were then evenly distributed into sterile polypropylene jars. A sterile plastic grid and a plastic pusher were placed in each jar to keep the vegetables submerged in the exudate once the lid was screwed on. Spontaneous fermentation was carried out by incubating the jars at 20°C for 28 days. At each sampling time point (days 0, 2, 7, 14, 21 and 28), three fermentation jars were sampled as biological replicates (Figure 1).

**Figure 1:**
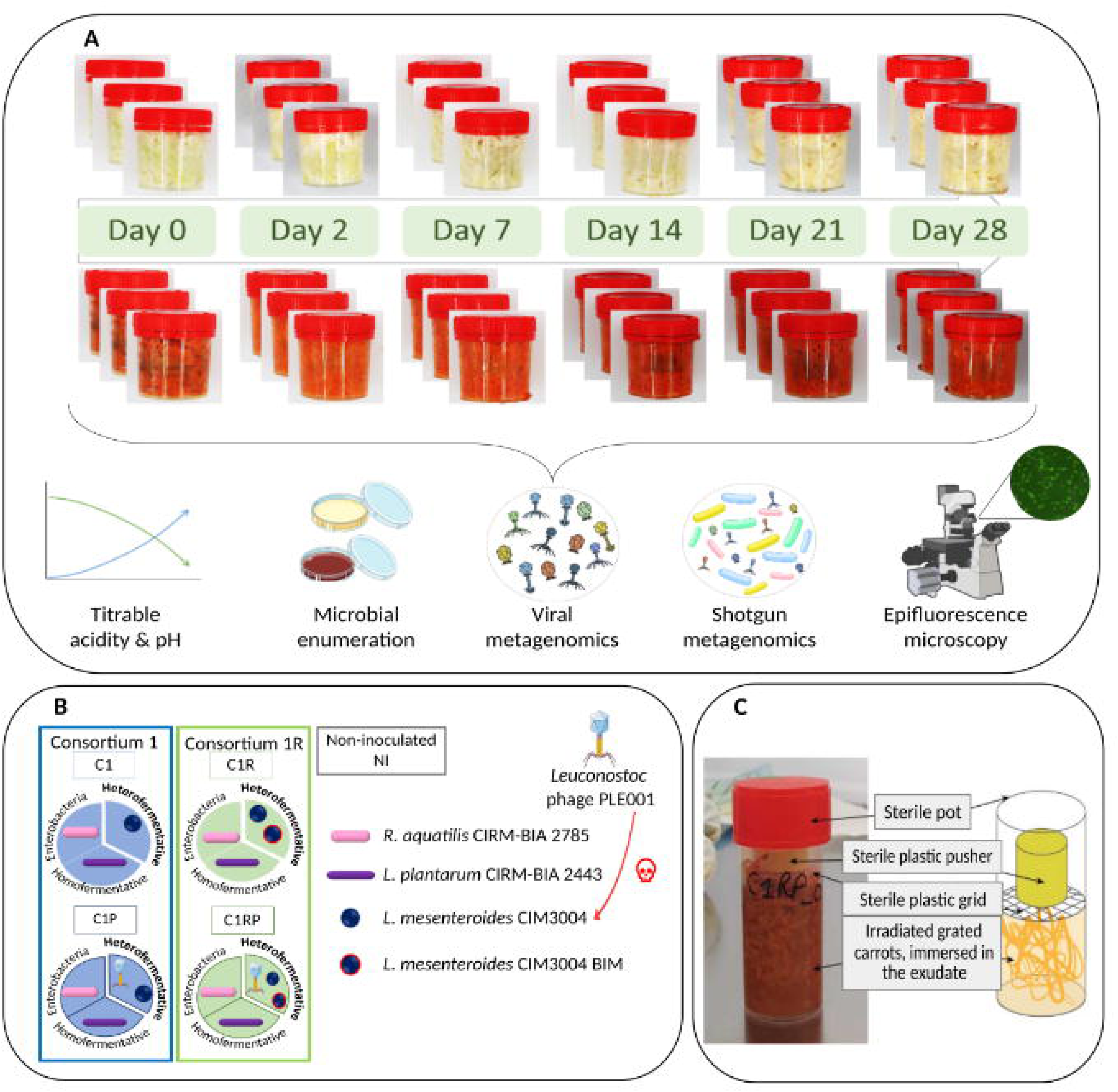
Schematic representation of the experimental design. A) The experimental design used to study the dynamics of microbial and viral communities. Cabbage and carrot were fermented in triplicate. All samples were analyzed by pH and titratable acidity measurements, microbial plate counts, shotgun metagenomics and viral metagenomics. (part of the illustrations used the NIAID NIH BIOART Source). B) & C) The experimental design using an ecological synthetic approach. B) Composition of the two consortia infected or not with phage PLE001. C) Illustration of the carrot fermentation model (25 g).

#### 3.1.2. Acidity measurement and microbial enumerations

For each sample, 2 g of product was placed in a cup for pH measurement using an InLab® Surface pH electrode (Mettler Toledo). Titratable acidity was measured on 5 mL of exudate using the HI931 Automatic Potentiometric Titrator (Hanna Instrument) after dilution to one-tenth with distilled water and 0.1 M NaOH (Merck) as the neutralizing solution. The titratable acidity was expressed in g of lactic acid/100 mL of exudate. For microbial counts, 10 g of product was homogenized with 90 mL of sterile saline solution (9 g/L NaCl) in a stomacher for 2 min at maximal speed (BagMixer®O400W, Interscience). Serial dilutions were surface-plated on specific agar media. Plate Count Agar (PCA) and Man-Rogosa-Sharpe agar (MRS, pH 6.5) (Biokar Diagnostics) were used for the determination of the total number of aerobic bacteria and LAB, respectively. Both media were supplemented with 22.5 mg/L amphotericin B to inhibit fungi. Fungal populations were enumerated using yeast-extract-glucose chloramphenicol (YEGC, Biokar Diagnostics). All incubations were carried out for 3 to 5 days at 28°C. MRS plates were incubated in anaerobic jars.

#### 3.1.3. Shotgun metagenomic analyses

For shotgun metagenomic analysis, 1 g of sample was diluted 10-fold with sterile saline solution (9 g/L NaCl), homogenized in a stomacher for 2 min at maximal speed (BagMixer®O400W, Interscience), and centrifuged at 10,000 x g for 30 min. The microbial pellet was washed with sterile saline solution, collected by centrifugation, and stored at −20°C for metagenomic DNA extraction. DNA extraction was performed as described previously (Cantuti Gendre et al., 2025). Briefly, the protocol included a pre-lysis step including Triton, lysozyme and mutanolysin. DNA was further extracted and purified using the DNeasy® PowerFood® kit (Qiagen). The vortexing step was replaced by two cycle of homogenization in a Precellys® Evolution homogenizer at 6500 rpm for 45 sec (Bertin Instruments). The DNA was quantified using the Qubit dsDNA HS Assay Kit (ThermoFisher Scientific). Shotgun sequencing was performed by Eurofins Genomics (Konstanz, Germany) using a NovaSeq platform (Illumina, San Diego, CA, USA). A minimum of 5 million paired-end reads (150 bp) were produced per sample. The taxonomic affiliation of the raw reads was carried out using Kaiju v.1.9.2 (Menzel et al., 2016) with nr_euk kaiju database (2023-05).

#### 3.1.4. Purification of the viral fraction and viral metagenomic analysis

The viral fraction was extracted from 10 g of fermented vegetables according to the protocol described in our previous study (Cantuti Gendre et al., 2025). Briefly, samples were diluted in Tris-HCl 10 mM, NaCl 200 mM buffer. Vegetable debris and microbial cells were removed using centrifugation and filtration (0.45 µm). Viral particles were precipitated using PEG 8000. Upon centrifugation, the pellet was resuspended in 1 mL of SM buffer (Tris-HCl 50 mM, NaCl 200 mM, MgSO4 10 mM) and stored at 4°C. DNA was recovered from viral particles following the procedure described in our previous work (Cantuti Gendre et al., 2025), comprising a DNase treatment to digest free DNA, capsid digestion with Proteinase K and SDS, and phenol-chloroform extraction. Viral ssDNA and dbDNA were amplified using the REPLI-g® Single Cell kit (Qiagen) following the manufacturer’s instructions. Amplified DNA was purified using AMPure® XP beads (Beckman Coulter) and sent to Eurofins Genomics (Konstanz, Germany) for shotgun sequencing using a NovaSeq platform (Illumina). A minimum of 5 million paired-end reads (150 bp) were produced per sample. Raw reads were decontaminated to remove those matching the cabbage and carrot genomes (accession numbers: GCA_000695525.1, GCA_001625215.1) according to the procedure detailed in Yeoh et al. (2021), and quality-filtered using Trimmomatic v0.39 (Bolger et al., 2014). Decontaminated and trimmed reads were subsampled to 100 k, 500 k, and 1000 k using seqtk version 1.3 (https://github.com/lh3/seqtk). All read sets were then independently assembled into contigs with SPAdes v3.15.3 (Bankevich et al., 2012), retaining those longer than 2,000 bp. Contigs from each assembly were clustered according to Shah et al. (2023). Briefly, pairwise alignments were performed using BLAT (Kent, 2002), and chimeric contigs (>110% self-alignment) were removed. Remaining contigs were clustered at 90% identity × coverage, with the longest contig in each cluster selected as the representative sequence. Cluster representative sequences were screened using VIBRANT v1.2.1 (Kieft et al., 2020), VirSorter2 v2.2.4 (Guo et al., 2021), and CheckV v0.8.1 (Nayfach et al., 2021). Clusters were retained as viral operational taxonomic units (vOTUs) if their representative sequence met at least one of the following criteria: classification as “complete”, “high”, or “medium” quality by VIBRANT or CheckV, or as “full” by VirSorter2. To estimate the relative abundance of each vOTU in samples, trimmed reads were mapped to vOTUs using bwa-mem2 v2.2.1 (Vasimuddin et al., 2019). Alignments were filtered with msamtools v1.1.3 (https://github.com/arumugamlab/msamtools) to retain only contigs detected with high confidence. First, msamtools filter was used with the option -l 80, -p 95, -q 80, --besthit. Then, msamtools coverage was used with the “--summary” option to estimate sequence coverage and only alignments with more than 50% coverage were kept to construct the final abundance table. vOTU host prediction was performed with iPHoP v1.2.0 (Roux et al., 2023). Similarity to known bacteriophages was assessed using BLASTn v.2.13.0 (Camacho et al., 2009) against the Core Nucleotide BLAST and ICTV databases. Unknown phages were annotated using Pharokka (Bouras et al., 2023). Analyses of the vOTUs abundance table, including calculation of the Shannon index, principal coordinate analysis (PCoA) based on Bray-Curtis dissimilarity, permutational analysis of variance (PERMANOVA), composition analysis, and heatmap visualization, were conducted in R using the following packages: Phyloseq v1.44.0 (McMurdie and Holmes, 2013), vegan v2.6-8 (Oksanen et al., 2024), pheatmap v1.0.12 (Raivo Kolde, 2010), ggplot2 v3.5.1 (Wickham, 2016), and tidyverse v2.0.0 (Wickham et al., 2019).

#### 3.1.5. Quantification of virus-like particles by epifluorescence microscopy

A volume of 100 μL of the viral fraction was 10-fold diluted in SM buffer and filtered through 0.02 μm Anodisc filters (Anodisc 25 mm diameter, Whatman). Each filter was then incubated on a 75 μL drop of 1X SYBR Gold (Invitrogen) in the dark for 15 min. After removal, filters were dried in the dark and mounted on a glass slide with Fluoromount-G (Invitrogen) and a coverslip. Microscopic observations were performed using a BX51 microscope (Olympus) equipped with a 100 × oil-immersion objective (Olympus U PlanFI(C1)Japan ×100/1.30 OILPh3∞/0,17). Ten images were captured per slide in GFP fluorescence channels (with an excitation filter FITC wavelength of 460-490 nm and emission filter wavelength of 520 nm). The images were captured using a DP23 Camera with the CellSens EVIDENT 2024 software (Olympus). The number of particles was determined using Icy software (de Chaumont et al., 2012) with the spot detector tool. Detection parameters were set to identify bright spots against a dark background with intensity ranging from 100 to 200, depending on the image. A filter was used to accept only objects with a size between 10 and 100 pixels. The final virus-like particles (VLP) concentration was calculated relative to the initial weight of the fermented vegetable sample, considering the ratio between the image area and the filter area, as well as the sample dilution factor. The detailed protocol is described in our previous article (Cantuti Gendre et al., 2025).

### 3.2. Synthetic microbial ecology experiment

#### 3.2.1. Bacterial strains, phage, and culture conditions

The synthetic community used in this study comprised three bacterial strains: Rahnella aquatilis CIRM-BIA2785 and Lactiplantibacillus plantarum CIRM-BIA2443 obtained from the CIRM-BIA collection (INRAE STLO, Rennes, France, https://collection-cirmbia.fr/), and Leuconostoc mesenteroides CIM 3004 obtained from the FranceMIL collection (CNIEL, France, https://www.francemil.fr/). Phage PLE001, which infects L. mesenteroides CIM 3004, was also obtained from the FranceMIL collection. R. aquatilis CIRM-BIA2785 was cultivated in a broth medium composed of tryptone (5 g/L), yeast extract (2.5 g/L) and D-Glucose (1 g/L) at 30 °C without agitation. L. plantarum CIRM-BIA2443 and L. mesenteroides CIM 3004 were cultivated in Man-Rogosa-Sharpe broth (MRS broth, Biokar Diagnostics) at 30 °C without agitation. Phage PLE001 propagation was carried out in MRS broth supplemented with 10 mM CaCl_2_. Briefly, a bacterial culture of L. mesenteroides CIM 3004 (OD_600nm_ = 0.2) was infected with 100 μL of a phage lysate obtained from the culture collection. The infected culture was incubated at 30 °C until lysis occurred. Cell debris were discarded by centrifugation at 4,700 × g for 20 min. The supernatant was filtered (0.45 μm PES syringe filters, Sartorius) and stored at 4 °C. The stock concentration was determined by phage titration using a classical double-layer plaque assay (Kropinski et al., 2009).

#### 3.2.2. Generation of a bacteriophage-insensitive mutant (BIM)

1. L. mesenteroides CIM 3004 was cultured in MRS broth supplemented with 10 mM CaCl_2_ at 30°C without agitation until the OD_600nm_ value reached 0.2. The culture was then infected with 1 μL of serial dilutions of the PLE001 phage stock (∼10 PFU/mL) and further incubated for 24 h. The culture infected with the 1:10-diluted PLE001 stock underwent complete lysis before subsequently regaining turbidity. This culture was then plated on MRS agar and incubated at 30°C under anaerobic conditions. A single colony was isolated and subsequently subcultured every 24 hours for seven consecutive passages in MRS broth supplemented with 10 mM CaCl_2_, with infection at each transfer using 1 μL of 1:10-diluted PLE001 stock. Cultures were incubated at 30°C under anaerobic conditions. After each transfer, the sensitivity of one isolated clone to phage PLE001 was evaluated using a spot assay. Briefly, serial dilutions of the PLE001 stock were prepared in SM buffer Then, 8 µL of each dilution were spotted onto a lawn containing L. mesenteroides CIM 3004, using the standard double-layer agar technique. After overnight incubation at 30°C, the phage titer was determined by counting lysis plaques at the lowest countable dilution. The final clone, designated as an insensitive mutant (CIM 3004 BIM), was selected after seven transfers and exhibited no lysis plaques at any phage dilution, indicating complete resistance to PLE001 (data not shown).

#### 3.2.3. Genome sequencing and variant calling

The reference genome of L. mesenteroides CIM 3004 was assembled using a combination of long- and short-read sequencing technologies. DNA was extracted using the NucleoMag HMW DNA kit (Macherey-Nagel) according to the manufacturer’s instructions, with an additional pre-lysis step as described below. The bacterial pellet was resuspended in 18 µL of pre-lysis buffer (20 mM Tris-HCl [pH 8], 2 mM EDTA, 1.2% Triton X-100, 20 mg/mL lysozyme). Ten units of mutanolysin, free of DNA contaminants (Sigma-Aldrich), were added, and the suspension was incubated for 35 min at 37 °C. After purification, DNA was sent to BMKGene (Münster, Germany) for sequencing using both a NovaSeq platform (Illumina, paired-end 150 bp, 100X coverage) and a Revio platform (PacBio, HiFi sequencing, 30X coverage). Genome assembly and annotation were carried out by BMKGene. Briefly, long reads from PacBio sequencing were assembled using Hifiasm v0.12 (Cheng et al., 2021) and circularized using Circlator v1.5.5 (Hunt et al., 2015). The sequence was further polished using Illumina short-reads with Pilon v1.22 (Walker et al., 2014). Coding DNA sequences were predicted using Prodigal v2.6.3 (Hyatt et al., 2010). For functional annotation, predicted proteins were compared to the nr, Swiss-Prot, TrEMBL (Bairoch and Apweiler, 1996), KEGG (Kyoto Encyclopedia of Genes and Genomes) (Kanehisa et al., 2004), and eggNOG (Powell et al., 2014) databases using BLASTP v2.2.29 (Altschul et al., 1990), with sequence identity >90% and coverage >90% of the target sequence as thresholds. Blast2GO v2.5 (Conesa et al., 2005) was used for Gene Ontology (GO) annotation, and HMMER v3.4 (Eddy, 2011) was used to compare the sequences against the Pfam database (Mistry et al., 2021).

Genome sequencing of the mutant (CIM 3004 BIM) was performed using short-read sequencing, and the resulting data were used for variant calling. DNA extraction was performed using the DNeasy® PowerFood® kit (Qiagen) according to the manufacturer’s instructions with slight modifications. Briefly, the protocol included the same pre-lysis step described above, and the vortexing step was replaced by homogenization using a Precellys® Evolution homogenizer (Bertin Instruments) for two 45 s mixing steps at 6,500 rpm. After purification, DNA was sent to BMKGene for sequencing using a NovaSeq platform (Illumina, paired-end 150 bp, 100X coverage). Comparative genomic and variant analysis were carried out by BMKGene. Briefly, raw sequencing reads were quality-filtered using FastP v1.0.0 (Chen, 2023) and mapped to the L. mesenteroides CIM 3004 reference genome using bwa-mem2 v2.2.1 (Vasimuddin et al., 2019). Mapped reads were sorted, and duplicate reads were removed using samtools v1.9 (Danecek et al., 2021). Single-nucleotide polymorphisms (SNPs) and insertions/deletions (InDels) were called using the HaplotypeCaller module in GATK v3.8 (McKenna et al., 2010) and filtered using the following parameters: QD < 2.0, MQ < 40.0, FS > 60.0, QUAL <30.0, MQrankSum < −12.5, ReadPosRankSum < −8.0, -clusterSize 2, -clusterWindowSize 5. SNP annotation was performed based on the reference genome using SnpEff v3.6c (Cingolani et al., 2012).

#### 3.2.4. Design of simplified consortia and laboratory-scale fermentations

Simplified microbial consortia were designed as described in Figure 1 and Table 1. They comprised R. aquatilis CIRM-BIA2785 and L. plantarum CIRM-BIA2443 and were expanded with the addition of L. mesenteroides CIM 3004 (consortium C1) or an equal mixture of L. mesenteroides CIM 3004 and its phage PLE001-resistant mutant derivative, L. mesenteroides CIM 3004 BIM (consortium C1R). C1 and C1R consortia were inoculated into sterile carrot matrices without phage or with the addition of phage PLE001 (C1P and C1RP). Initial CFUs and PFUs counts are provided in Table 1. A non-inoculated assay (NI), with no exogenous addition of bacteria and phage, was included as a control.

**Table 1:**
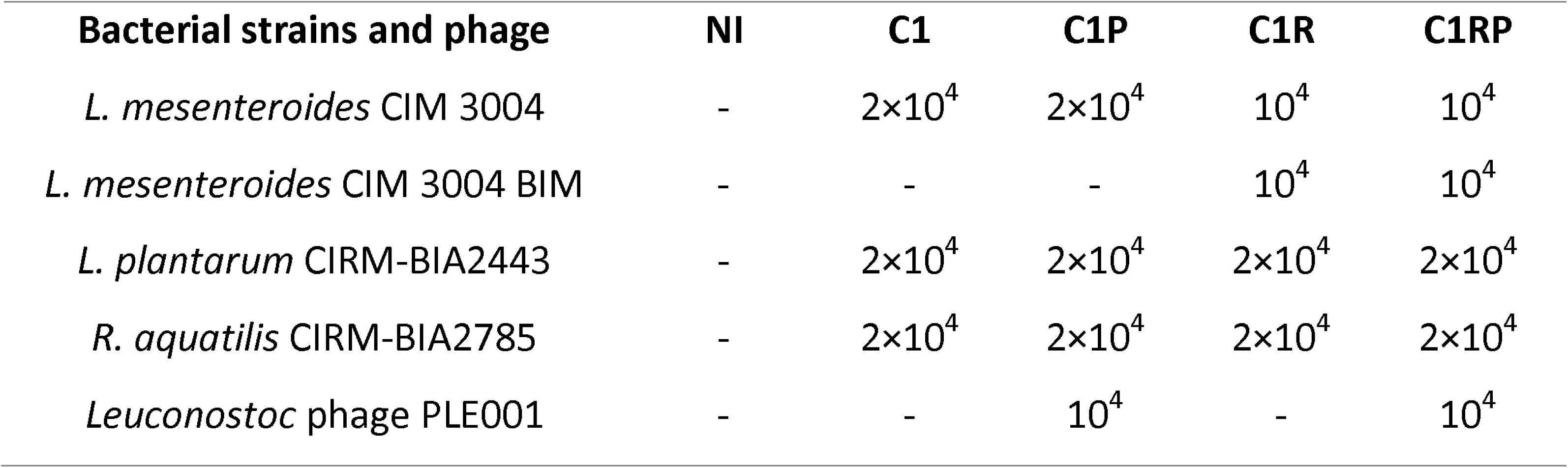
Concentrations of bacterial strains (CFU/g) and PLE001 phage (PFU/g) used for inoculation in the model system of carrot fermentation.

Fresh organic carrots were grated (size approx. 2 mm), conditioned, and vacuum-packed in plastic bags (Cenpac). They were decontaminated by irradiation with 10 meV electron beams (Rhodotron TT300, IBA) yielding a fresh dose of 8 kGy, and stored at −40°C until use. Carrots were thawed for 24 hour at 4°C before each batch production. On the production day (day 0), 700 g of grated carrots was mixed with sterile sea salt (1.5% w/w) in a sterile beaker placed under a biological safety cabinet, homogenized manually using sterile gloves. The mixture was incubated for 1 hour at room temperature. To ensure an even distribution of vegetable and exudate among all jars, we followed the procedure described in the spontaneous fermentation experiment, except that the exudate was separated into five sterile flasks for inoculation (concentrations in Table 1) prior to distribution into the jars. Both the vegetable and the inoculated exudate were equally distributed among the different jars to obtain a final product weight of 25 g per jar. A sterile plastic grid and a plastic pusher were placed in each jar to ensure that the vegetable remained submerged in the exudate after screwing of the lid (Figure 1C). The jars were incubated at 19°C for 28 days. Four independent batches of fermented carrots were produced on different dates and used as biological independent replicates.

#### 3.2.5. Sample collection and analysis

Samples were collected at three different time points during fermentation (days 0, 3 and 28). On each sampling day, one jar per condition (NI, C1, C1P, C1R, C1RP) was opened under sterile conditions. Samples were processed as described above for CFU and PFU enumerations, pH and titratable acidity measurements. In addition, lactic acid concentration was determined from 1.5 mL of exudate using the D-/L-Lactic Acid (D-/L-Lactate) Rapid Assay Kit (K-DLATE, Megazyme) according to the manufacturer’s instructions.

## 4. Results

### 4.1. Monitoring of laboratory scale models mimicking spontaneous cabbage and carrot fermentation

Spontaneous fermentations of cabbage and carrot were conducted at laboratory scale. To ensure that these laboratory-scale fermentations were representative of actual production systems, the process was monitored over the entire fermentation process (28 days) by measuring pH, titratable acidity, viable counts of total bacteria, LAB, and yeasts, as well as counts of VLP (Figure 2).

**Figure 2:**
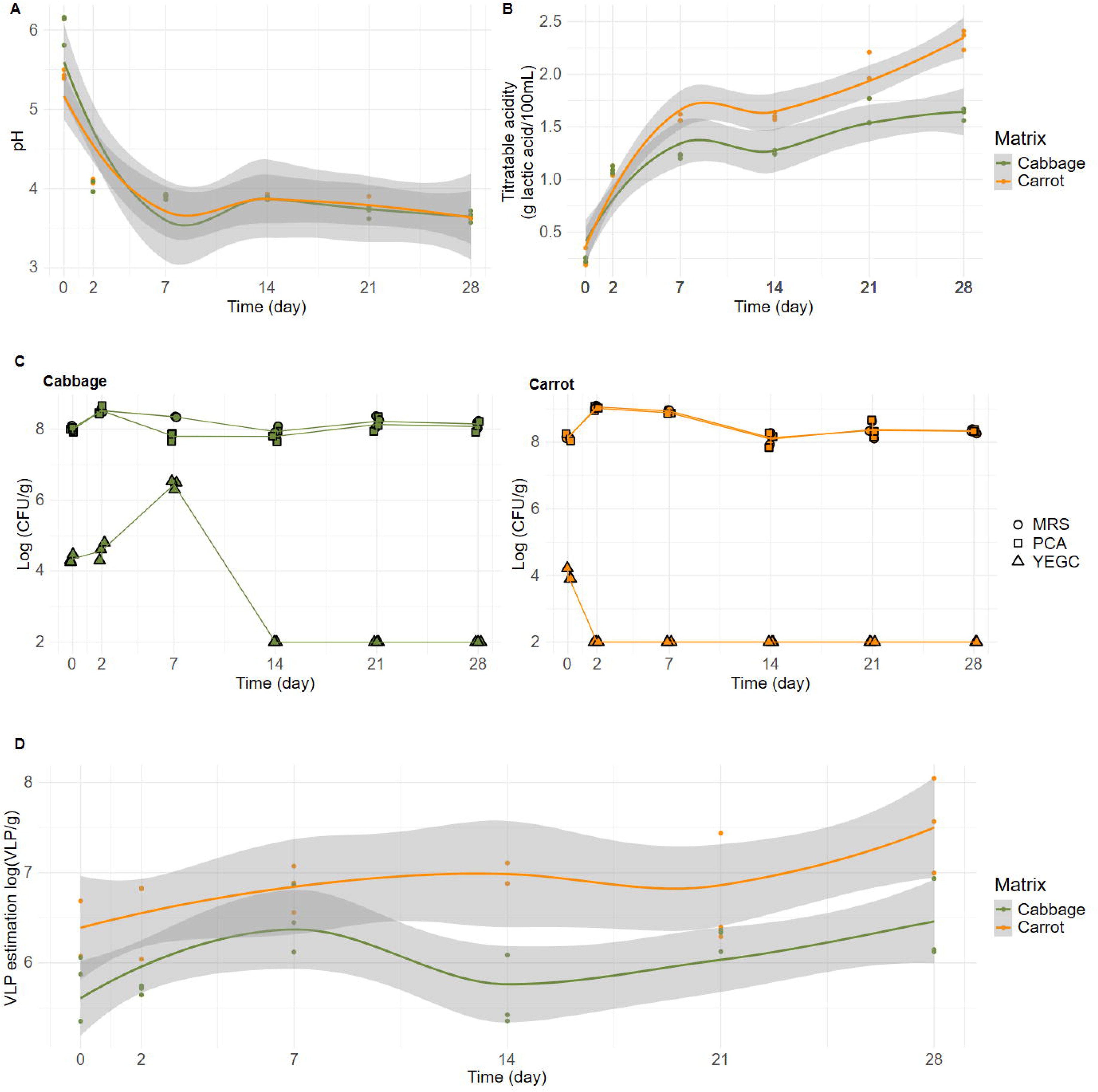
Dynamics of key indicators during the fermentation of vegetables. A) pH. B) Titratable acidity expressed in g of lactic acid per 100 mL of exudate. C) Viable counts of total bacteria, LAB and yeasts as determined by plating on PCA, MRS and YEGC media, respectively. D) Estimation of virus-like particle concentration using epifluorescence microscopy. Three fermentation batches were produced for each vegetable and used as biological replicates. Each dot represents the value obtained for one replicate. Samples were collected through the fermentation process on days 0, 2, 7, 14, 21, and 28. Solid lines correspond to the average values, and the gray envelopes indicate the confidence interval at 95%.

A decrease in pH was observed during both cabbage and carrot fermentations, and similar final values of 3.7 (± 0.1) and 3.6 (± 0.1) were reached after 28 days, respectively (Figure 2A). Acidification was rapid and the pH dropped below 4.6 after two days. This value is generally considered as a threshold from a food safety perspective (Buckenhueskes, 2015; Snyder et al., 2020). Titratable acidity, which reflects the concentration of acids (mainly lactic acid) in the exudate, increased from 0.2 (± 0.02) to 1.6 (± 0.1) g/100 mL for cabbage and from 0.3 (± 0.1) to 2.3 (± 0.1) g/100 mL for carrots (Figure 2B). An acidity titer of 1 g/100 mL is the minimal threshold for considering a successful fermentation by French sauerkraut producers (INAO, 2018).

Both fermented vegetables had high initial counts of LAB, reaching up to 8 (± 0.1) log CFU/g (Figure 2C). LAB population slightly increased on day 2, progressively declined thereafter until day 14, and stabilized at 7.9 (± 0.1) and 8.1 (± 0.1) log CFU/g until the end of fermentation for cabbage and carrot, respectively. This dynamic is consistent with previous studies on similar products (Lu et al., 2003; Wuyts et al., 2018). A similar trend was observed for total aerobic bacteria. Yeasts were present at 4.3 (± 0.1) and 4.0 (± 0.2) log CFU/g at the initial stage of fermentation in cabbage and carrot, respectively. However, they were no longer detected after day 2 in carrot and day 7 in cabbage.

The concentration of virus-like particles, as estimated by epifluorescence microscopy, increased during fermentation for both vegetables, from 5.8 (± 0.4) to 6.4 (± 0.5) log particles/g of cabbage, and from 6.4 (± 0.4) to 7.5 (± 0.5) log particles/g for carrot (Figure 2D). These values fall within the same range as those obtained in our previous work on fermented vegetables (Cantuti Gendre et al., 2025).

Taken together, these results indicate that our laboratory-scale fermentations exhibit characteristics comparable to conventional fermentations and can therefore be used as reliable model systems for our study. Moreover, these models offer the important advantage of enabling kinetic investigations throughout the fermentation process, which is not feasible in current industrial settings.

### 4.2. Succession of phage populations reflects the dynamics of the bacterial community during spontaneous fermentation of vegetables

We further analyzed the same samples to explore the dynamics of both viral and microbial communities during fermentation. Total microbial and viral DNA were extracted from 18 carrot and 18 cabbage kinetic samples (triplicate samples for each of the 6 time points) and subjected to shotgun metagenomic and viral metagenomic analysis, respectively.

First, the overall microbial composition of the fermented vegetables was assessed by taxonomic affiliation of the metagenomic reads using Kaiju (Menzel et al., 2016). Half of the reads (53.96%) were successfully assigned, with the vast majority corresponding to bacteria (92.5% −99.7%). Eukaryotic and viral reads were also detected but in smaller proportions, between 0.03% and 4.5% and 2.2% and 6.9%, respectively (Supplementary Figure S2). Lactobacillaceae was the most dominant bacterial family in our dataset, with Leuconostoc as the main genus in fermented carrots (Figure 3A). In cabbage, this genus was accompanied by other LAB genera including Secundilactobacillus, Lactiplantibacillus, Levilactobacillus, Pediococcus and Weissella. In addition, some genera belonging to the Pseudomonadota, including Rahnella, Enterobacter and Pantoea, were detected at low proportions in our dataset.

**Figure 3:**
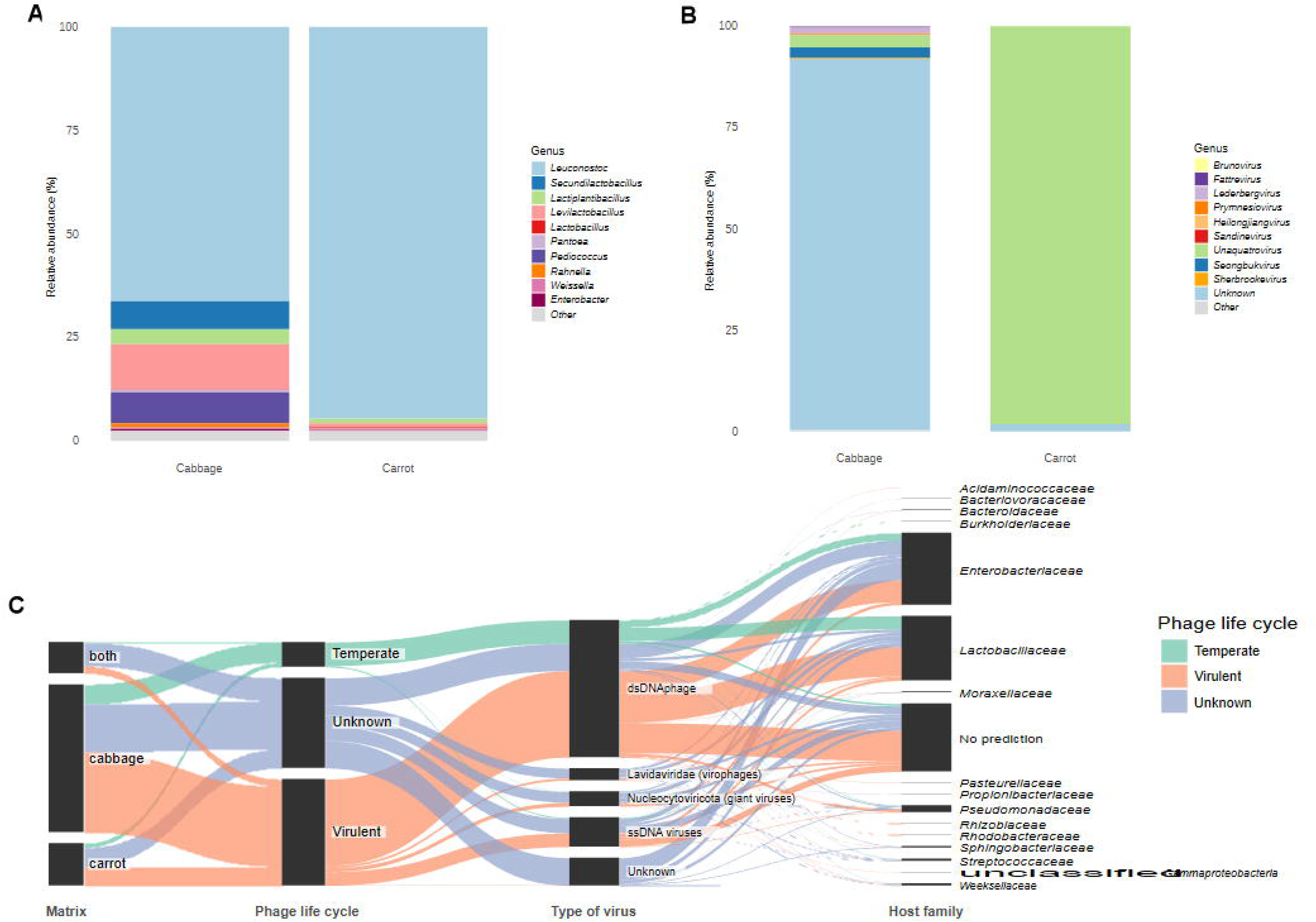
Composition of the microbial community of spontaneous fermented cabbage and carrot. Barplots of bacterial (A) and viral (B) communities. Relative abundance of reads (%) at the genus level. C) Parallel plot of the viral composition of fermented cabbage and carrot. Each vOTU is represented according to the vegetable in which it is present, its predicted life cycle (based on Vibrant prediction), virus type (according to VirSorter2 prediction), and host family (based on iPHoP prediction).

The same global approach was applied to the reads obtained from free viral particles through a viral metagenomic approach. In total, 794 species-level viral operational taxonomic units (vOTUs) were detected in our samples. Following taxonomic affiliation of the vOTUs by PhaBOX2 (Shang et al., 2023), it was found that cabbage samples contained many unknown vOTUs, while the majority of vOTUs detected during carrot fermentation belonged to the Unaquatrovirus genus (Figure 3B). The majority of the vOTUs present in our dataset (530) were unique to cabbage samples, while 152 were specifically detected in carrots. Only 112 vOTUs were shared between the two types of fermented vegetables (Figure 3C). The vOTUs corresponded mainly to dsDNA viruses. They were predicted to infect bacteria from the Enterobacteriaceae (258 vOTUs) and Lactobacillaceae (232 vOTUs) families, according to VirSorter2 (Guo et al., 2021) and iPHoP (Roux et al., 2023). Only a few of them (89 vOTUs) were predicted as temperate phages by VIBRANT (Kieft et al., 2020).

Once the overall composition of the microbial and viral communities was characterized, we next evaluated the dynamics of the different populations during fermentation using our shotgun and viral metagenomic datasets. Because our dataset includes two independent case studies, namely carrot and cabbage fermentations, we analyzed each of them separately to account for their specific microbial and viral trajectories.

In cabbage (Figure 4), a rapid and substantial change occurred within the first two days of fermentation. Taxonomic affiliation of the metagenomic reads (Figure 4A) revealed that the bacterial community was dominated by members of the Leuconostoc genus at day 0 (81.5% to 88.3% of the reads), together with several genera from the Pseudomonadota phylum, including Rahnella (3.9% to 5.1%), Pantoea (2.1% to 4.7%) and Enterobacter (1.2% to 3.4%). At day 2, the proportion of reads corresponding to Pseudomonadota decreased, and Leuconostoc dominated the ecosystem. A major shift occurred after day 2, with the successive emrgence of other LAB genera, including Pediococcus and Secundilactobacillus between day 7 and day 14, and Levilactobacillus and Lactiplantibacillus after day 14. The strong effect of the fermentation stage on the microbial community structure was confirmed by PCoA analysis based on Bray-Curtis dissimilarity, and statistically supported by a PERMANOVA test (p = 0.001, R² = 0.897, Supplementary figure S3A). Regarding viruses, analysis of the vOTUs abundance table also revealed the succession of different populations during the fermentation process (Figure 4B), which was supported by PCoA analysis based on Bray-Curtis dissimilarity and PERMANOVA test (p = 0.001, R =0.691, Supplementary figure S4A). Six vOTUs clusters were detected according to the gap statistics method (Tibshirani et al., 2001) and distributed across four fermentation stages, as observed for bacteria. At day 0, a large cluster (Cluster III) comprising vOTUs predicted to infect various bacteria belonging to the Pseudomonadota phylum was detected. At day 2, cluster II and VI, both comprising vOTUs infecting members of the Leuconostoc genus became dominant, while Cluster III disappeared. At day 7, cluster V was the most dominant vOTU cluster and comprised phages predicted to infect Pediococcus. Finally, clusters I and IV, detected at low abundance at the early stages of fermentation, became more abundant after day 7. These clusters were primarily composed of vOTUs predicted to infect the bacterial genera: Levilactobacillus,

**Figure 4:**
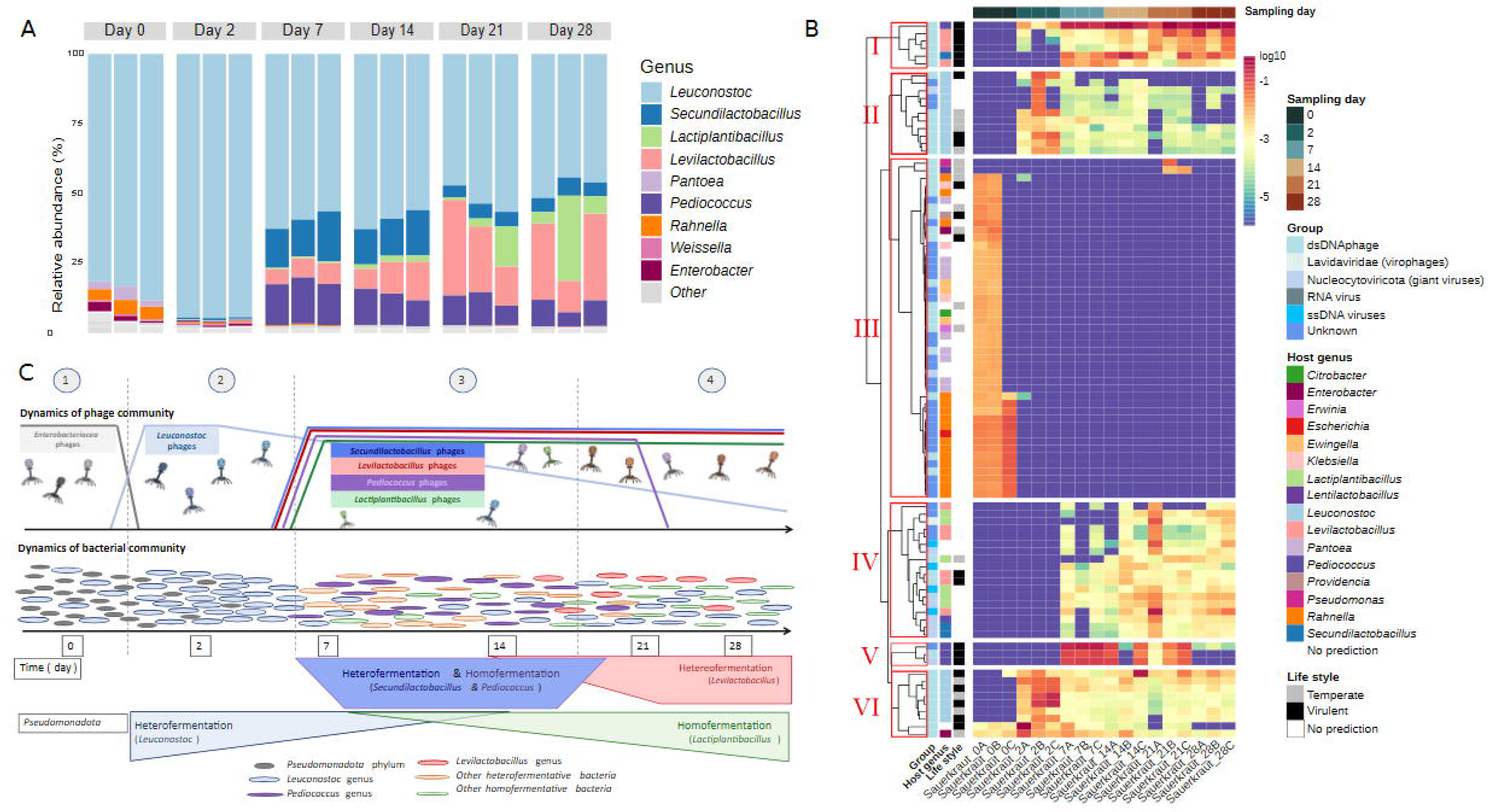
Temporal variation of bacterial and viral communities during cabbage fermentation. A) Temporal variation of bacterial community. Relative abundance of reads per sample (%) at the genus level. B) Distribution of the most abundant vOTUs during cabbage fermentation. The colors on the heatmap represent the log-transformed relative abundance and range from blue (low relative abundance) to red (high relative abundance). Viral group, host genus, and lifestyle of the different vOTUs are indicated when available. C) Schematic representation of phage-bacteria dynamics during cabbage fermentation.

Lactiplantibacillus, and Secundilactobacillus. We finally applied a Mantel test to evaluate the statistical relationships between the Bray-Curtis distance matrices produced from the virome and bacteriome datasets. We observed a positive correlation (p = 0.001; r = 0.6086; 999 permutations), indicating that changes observed in the phage community structure are associated with bacterial successions during cabbage fermentation. To summarize, our data collected throughout the cabbage fermentation indicate that four bacterial groups developed successively, along with their associated phages (Figure 4C).

We next analyzed the carrot fermentation dataset to assess whether similar dynamics would occur. The vast majority of the metagenomic reads (92.5% to 98.4%) corresponded to LAB, and a lower diversity was observed as compared to cabbage samples. The genus Leuconostoc was dominant throughout the fermentation process (Figure 5A). Lactiplantibacillus and Levilactobacillus were sub-dominant populations in the early stages of fermentation, and increased after day 21. The effect of the fermentation stage on the composition of the bacterial community was evaluated by computing Bray-Curtis dissimilarity and principal coordinate analysis, and confirmed by a PERMANOVA test (p = 0.002, r2 = 0.808) (Supplementary figure S3B).

**Figure 5:**
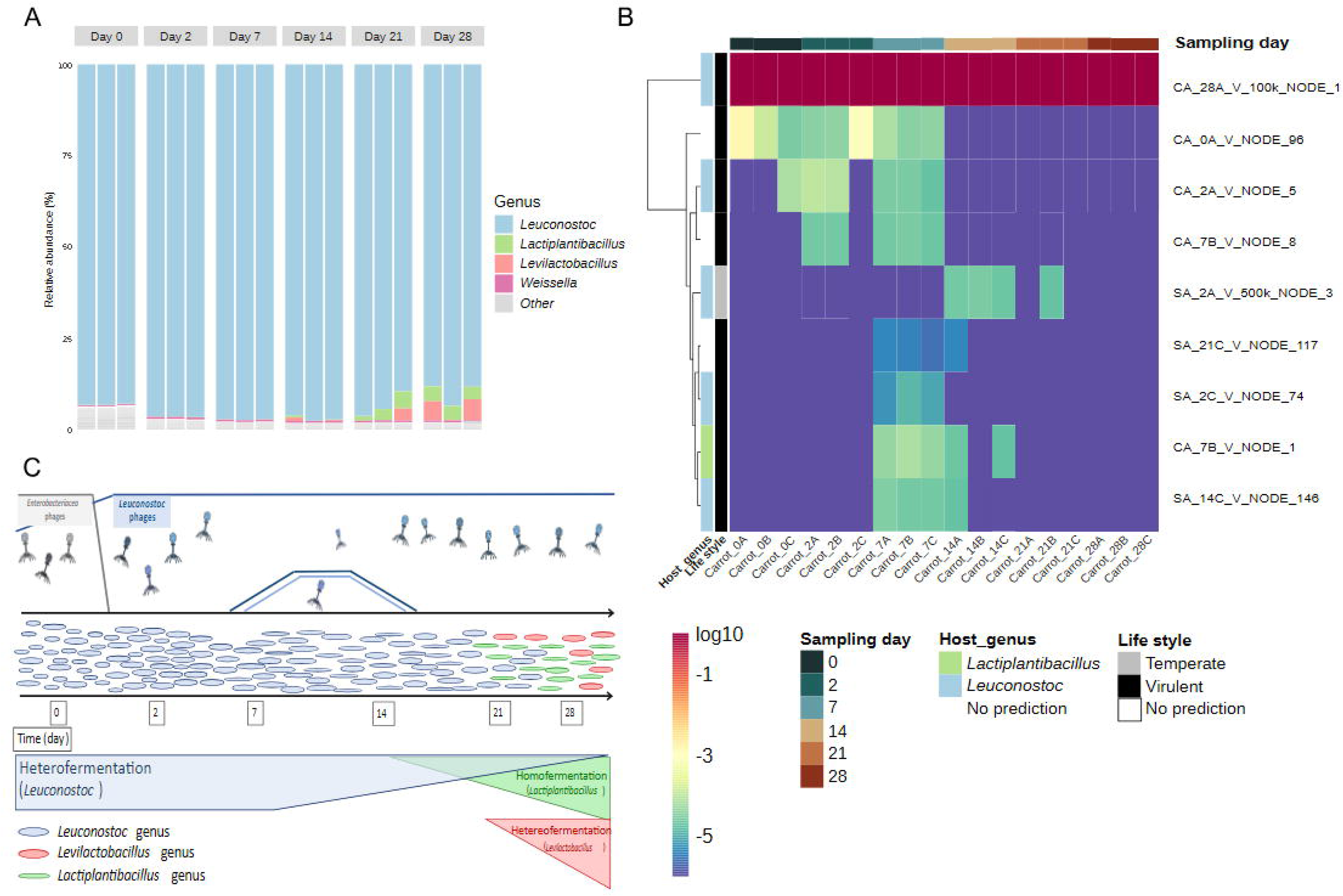
Temporal variation of bacterial and viral communities during carrot fermentation. A) Temporal variation of bacterial community. Relative abundance of reads per sample (%) at the genus level. B) Temporal variation of the virome during carrot fermentation. For better clarity, only vOTUs consistently detected across all three replicates were represented. The colors scale represents the log-transformed relative abundance, ranging from blue (low relative abundance) to red (high relative abundance). Viral group, host genus, and lifestyle of the different vOTUs are indicated when available. C) Schematic representation of phage-bacteria dynamics during carrot fermentation.

The viral diversity observed in carrot samples was also lower than in cabbage, as visualized through the heatmap of the most important vOTUs (Figure 5B). To assess potential variations in the viral community during carrot fermentation, Bray-Curtis dissimilarity was calculated, followed by principal coordinate analysis (Supplementary Data S3). The PERMANOVA indicated that the composition did not change significantly during fermentation (p > 0.05). The most abundant vOTU (CA_28A_V_100k_NODE_1) was predicted as a virulent Leuconostoc phage. Its relative abundance ranged from 98.7% of the reads at day 0 to 100% at day 28 (average value: 99.47%). This vOTU was 29 kb in size, and showed high sequence homology with the virulent phage 1-A4 infecting L. mesenteroides (93.82% identity; 84% coverage). This phage has been isolated from industrial sauerkraut in the USA and belongs to the Unaquatrovirus uv1A4 species (Lu et al., 2010) (Lu et al., 2010). Other vOTUs were detected at lower relative abundances (Figure 5B). Among them, six were successfully assigned to a bacterial host by iPHoP: five to Leuconostoc and one to Lactiplantibacillus. The later was detected between days 7 and 14. As summarized in Figure 5C, our results indicate that L. mesenteroides and its phage strongly dominated laboratory-scale fermented carrots throughout the fermentation process. Despite the low bacterial diversity observed, a clear succession of sub-dominant bacterial populations occurred during fermentation, accompanied by changes in the minor phage populations.

Taken together, our longitudinal data collected on both cabbage and carrot spontaneous fermentations demonstrated that viral population succession takes place during fermentation and follows the dynamics of the bacterial community. It therefore provides indirect evidence that multiple phage infections occur during vegetable fermentation. As the fermentation of carrots is a simpler process than that of cabbage, we decided to use carrots as a model for vegetable fermentation in our further investigations.

### 4.3. A single virulent phage can actively replicate and regulate the population level of its bacterial host during vegetable fermentation

We next applied a synthetic microbial ecology approach to provide direct evidence of active phage replication during fermentation. Based on the bacterial composition observed previously (Figures 4A and 5A), we identified the main bacterial genera that developed successively during spontaneous vegetable fermentation: the Enterobacteriaceae genus Rahnella at the beginning of the fermentation, followed by the genus Leuconostoc during the heterofermentative phase, and finally an increase in the genus Lactiplantibacillus during the homofermentative phase. We therefore developed a model system mimicking carrot fermentation (see Materials and Methods section), which involved inoculating sterilized carrots with a simplified community composed of one Enterobacteriacceae (Rahnella aquatilis CIRM-BIA2785) and two LAB (Leuconostoc mesenteroides CIM 3004 and Lactiplantibacillus plantarum CIRM-BIA2443) (Figure 1). This consortium (control condition C1) was then exposed to the virulent phage PLE001 (condition C1P) infecting L. mesenteroides CIM 3004. We also generated a bacteriophage-insensitive mutant (BIM) of CIM 3004 for phage PLE001 in order to mimic the emergence of resistant bacteria in the system (conditions C1R and C1RP). This mutant was later named CIM 3004 BIM.

To identify potential mutations associated with the resistance phenotype, we sequenced the genomes of both the parental strain (CIM 3004) and CIM 3004 BIM. This analysis showed that both plasmids from strain CIM3004 (30.8 kb and 44.5 kb) were present in the BIM derivative, while co-evolution with the phage resulted in the accumulation of six SNP all located in coding regions (Table 2). Among these mutations, one non-synonymous single nucleotide polymorphism (SNP) occurred in gene GE001349, which encodes a putative polysaccharide chain length regulator containing a Wzz/FepE/Etk N-terminal domain. This protein is the flippase component of a putative Wzx/Wzy-dependent pathway which is the primary intracellular mechanism used by LAB to synthesize heteropolysaccharides. Wzz dictates the specific chain-length distribution of the secreted exopolysaccharides (Islam and Lam, 2014). Surface polysaccharides serve as the primary receptors for bacteriophage adsorption in LAB (Mahony, 2021; Giesbers et al., 2023). The mutation observed in wzz may therefore alter the synthesis of surface polysaccharides, potentially reducing phage PLE001 adsorption efficiency and conferring resistance in CIM 3004 BIM.

**Table 2:**
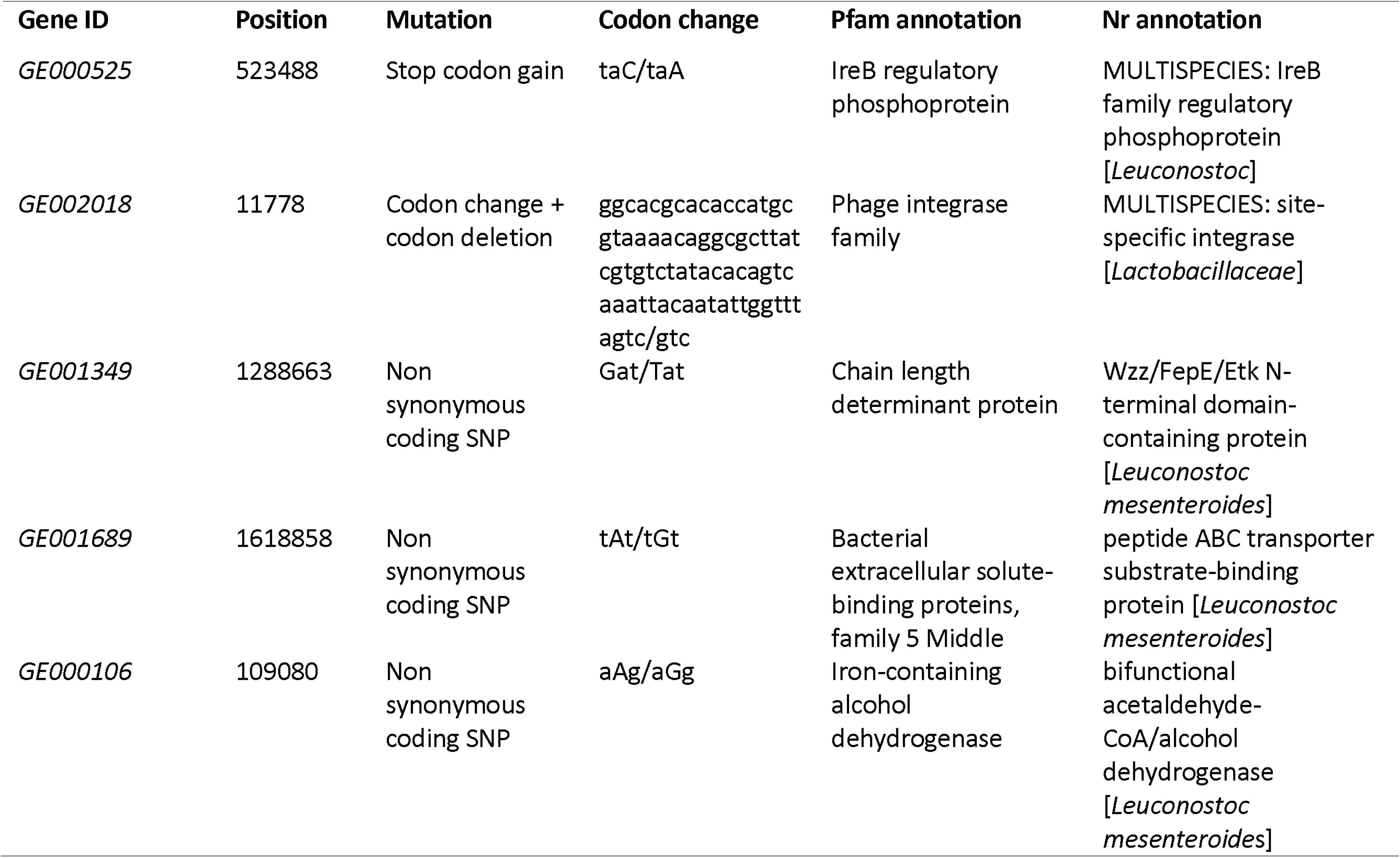
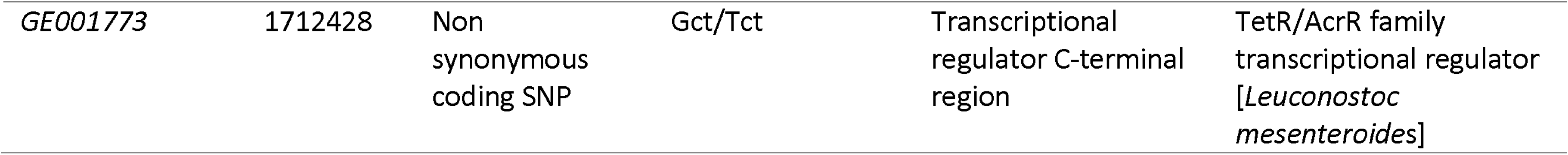
Annotation list of mutated genes of L. mesenteroides CIM3004 BIM.

We first examined whether our synthetic model of fermentation behaved as expected from a physico-chemical point of view, by monitoring pH and titratable acidity at three time points during fermentation (Figure 6). All tested conditions promoted rapid acidification of the medium compared to the non-inoculated samples. The final pH ranged from 3.5 ± 0.1 to 3.6 ± 0.1 (Figure 6A), and titratable acidity ranged between 1.6 ± 0.1 and 1.6 ± 0.1 (Figure 6B). No significant effect of phage PLE001 addition on pH or titratable acidity values was observed (Wilcoxon test, p > 0.05), indicating that the presence of phage PLE001 did not influence the global acidification process under the experimental conditions tested. We next analyzed lactic acid production (Supplementary Figure S5A). At day 0, the initial lactic acid concentrations were below or near the detection limits of the enzymatic kit. Lactic acid was produced across all consortia, reaching final concentrations of 15.7 ± 0.8 g/L to 17.1 ± 1.9 g/L. Moreover, the addition of phage PLE001 did not significantly affect lactic acid production in all consortia (Wilcoxon test, p > 0.05).

**Figure 6:**
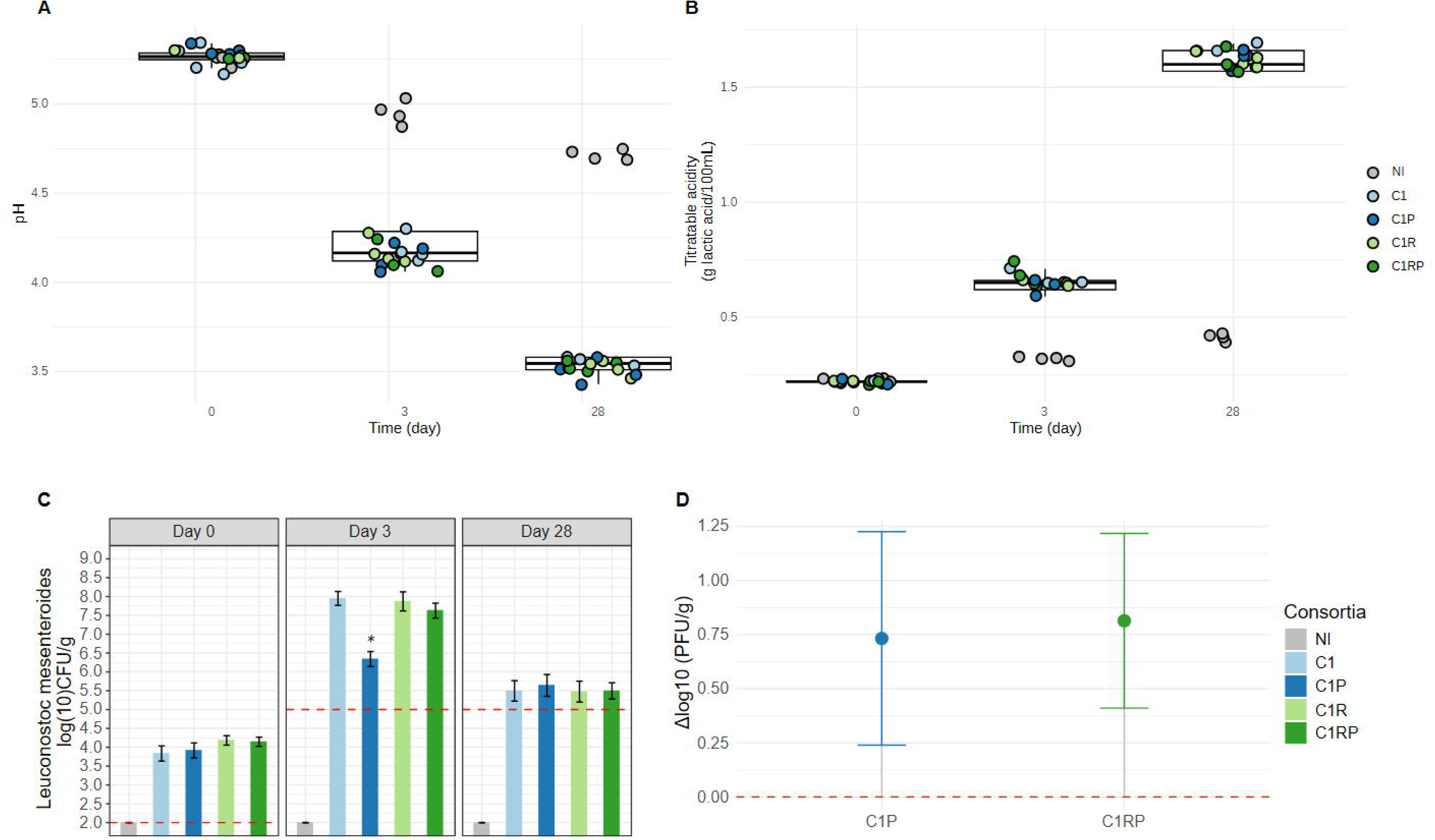
Kinetics of acidification and changes in the microbial and viral load during synthetic carrot fermentation. Boxplots representing the distribution of pH (A) and titratable acidity (B) values of each replicate for each of the four tested consortia and the control. Titratable acidity is expressed in g of lactic acid equivalent per 100 mL of exudate. C) Viable counts of Leuconostoc mesenteroides. Bar indicates the average value of four replicates (independent laboratory-scale productions) in log CFU/g of vegetables, with error bars representing the standard deviation. The dashed red line indicates the detection limit for microbial enumeration (2 CFU/g for day 0 and 5 CFU/g for day 3 and 28). D) Phage production dynamics between D0 and D3 of fermentation in consortia infected with phage PLE001. The Δlog10 represents the difference between PFU counts at D0 and D3.

We next investigated whether the presence of the Leuconostoc phage influenced microbial dynamics throughout the fermentation process using plate counting (Figure 6 and Supplementary Figure S5). First, no bacterial counts were observed in non-inoculated samples, indicating that no external contamination occurred in our experimental setup. In the absence of exogenously provided phage, we observed a rapid decline of Enterobacteriaceae (R. aquatilis CIRM-BIA 2785) (Supplementary Figure S5B), followed by the successive development of heterofermentative LAB (L. mesenteroides) and homofermentative LAB (L. plantarum CIRM-BIA 2443) (Figure 6 and Supplementary Figure S5C), which dominated at the end of fermentation. Our experimental design therefore reproduced the classical dynamics of bacterial populations observed earlier in spontaneous fermented vegetables. In the infected consortia (C1P and C1RP), phage PLE001 was detected at day 0 at a concentration of 4.0 ± 0.2 log PFU/g. This roughly corresponded to the targeted inoculum and a MOI of 1 (Supplementary Figure S5D). A statistically significant effect of phage addition on bacterial population levels was observed on day 3. The viable counts of Leuconostoc mesenteroides were significantly lower in consortium C1P than in C1 (Wilcoxon test, p < 0.05) (Figure 6C). This effect was accompanied by an increase in phage titer, reaching 4.7 ± 0.4 log PFU/g in condition C1P (Figure 6D). In contrast, similar CFU values were obtained in consortia C1R and C1RP, comprising both CIM 3004 (susceptible strain) and CIM 3004 BIM (insensitive mutant), while PFUs slightly increased in C1RP to 4.8 ± 0.3 log PFU/g on day 3, as observed in condition C1P.

Collectively, these results demonstrate that phage PLE001 actively replicates during fermentation whenever the susceptible strain is present, highlighting the potential influence of a virulent phage on the population level of its bacterial host.

## 5. Discussion

The dynamics of microbial communities in fermented vegetables have been extensively studied (Buckenhueskes, 2015; Pederson, 1969; Thierry et al., 2023). Current knowledge suggests that the initial microbiota associated with the raw material, typically dominated by Pseudomonadota and yeasts, is rapidly outcompeted by heterofermentative LAB, which are subsequently replaced by homofermentative LAB as fermentation progresses. Using shotgun metagenomics data, we observed similar dynamics in our laboratory-scale models of spontaneous fermentation. In the case of cabbage, our research confirms that the initial phase includes the rapid colonization of the raw material by Leuconostoc. Yet it is followed by the sequential progression of Secundilactobacillus, Pediococcus, and Levilactobacillus, before the subsequent increase of Lactiplantibacillus towards the end of fermentation. In carrots, Leuconostoc dominated the system throughout the fermentation process, but subdominant bacterial genera such as Levilactobacillus and Lactiplantibacillus were progressively observed after two weeks of fermentation.

Bacteriophages are frequently suspected to influence microbial assembly in fermented foods (Paillet and Dugat-Bony, 2021; Zhang et al., 2025), but experimental evidence is still largely lacking. Here, our viral metagenomics data highlighted that phage particles targeting all the key bacterial species are present in our experimental setup, confirming previous findings on commercial products (Cantuti Gendre et al., 2025). Furthermore, in both cabbage and carrot models of spontaneous fermentation, the observed successions of viral populations mirror the dynamics of the bacterial community. This observation supports the occurrence of active viral replication during fermentation, with population dynamics consistent with the Kill-the-Winner ecological model (Thingstad, 2000). The scenario describes a stabilizing frequency-dependent selection. Hence, when a particular microbial species (the “winner”) becomes dominant due to favorable conditions or resource availability, phages targeting that species proliferate, thereby controlling its population. In this scenario, virulent phages predominantly replicate in fast-growing bacteria and thereby suppress the competitive exclusion of slower-growing bacteria.

One limitation of studying phage-host interactions in natural communities using metagenomic approaches is that conventional MAG reconstruction, particularly from short-read data, often provides limited resolution of intraspecific diversity. This limitation is relevant even in a relatively simple ecosystem such as fermented carrots, because many phages have a narrow host range and may infect only a subset of strains within a bacterial species (Holtappels et al., 2023). The simplified community used for carrot fermentation in our synthetic ecology experiment, provided a tractable experimental framework for moving beyond the correlative patterns inferred from metagenomic time series obtained from natural communities and directly assessing the ecological consequences of active phage replication during fermentation. Despite its reduced complexity, the model reproduced the main ecological and functional features of vegetable fermentation, such as succession of three bacterial populations and rapid acidification of the raw material. The introduction of phage PLE001 altered the population dynamics of the susceptible strain L. mesenteroides CIM 3004 demonstrating active phage replication, but without preventing completion of the fermentation. This finding highlights a potential decoupling between taxonomic dynamics and ecosystem functioning, as substantial perturbation of an individual bacterial population does not necessarily translate into loss of the community-level function to which that population contributes. In this case, the reduced contribution of the susceptible heterofermentative population was apparently mitigated by the subsequent activity of the homofermentative LAB (L. plantarum CIRM-BIA2443), suggesting that functional redundancy and microbial succession can confer resilience to phage predation in this ecosystem. Similar observations were described in complex dairy starter cultures, in which strain diversity and the emergence of phage-insensitive populations (e.g. resistant clones) help maintain acidification despite phage pressure (Erkus et al., 2013; Spus et al., 2023). Although more diverse natural communities may exhibit additional direct and indirect responses, our results support the view that phages can redirect microbial succession without necessarily compromising fermentation performance.

Controlled experiments conducted in other microbial ecosystems indicate that such community-wide effects of phage infection are common in nature. For example, (Bouvier and Giorgio, 2007) incubated marine bacterial communities in virus-depleted seawater and observed the striking emergence of bacterial taxa that were normally rare in situ but appeared particularly effective at resource exploitation. The authors interpreted this pattern of selective mortality as a density-independent form of the Kill-the-Winner interaction, providing continuous background regulation of bacterial community composition alongside the density-dependent control of episodically abundant populations. In soil microcosms, experimental increases in phage pressure altered bacterial community composition and diversity and, in specific treatments, substantially increased soil ammonium concentrations, providing evidence that phages can influence both microbial community assembly and ecosystem functioning (Braga et al., 2020). Using germ-free mice colonized with a defined ten-member bacterial community, (Hsu et al., 2019) showed that the effects of species-specific phage infection propagated beyond susceptible hosts through the network of microbial interactions. These cascading effects altered non-targeted populations and produced measurable changes in the gut metabolome, demonstrating that even highly specific phages can have community-wide consequences for microbiome composition and functioning.

Together, these experimental studies show that active phage replication can affect not only susceptible hosts, but also community assembly and ecosystem functions. Such effects can arise mechanisms extending the direct lysis of susceptible cells (Burmeister et al., 2019; Paillet and Dugat-Bony, 2021; Brown et al., 2022). Phage activity can rapidly reduce dominant bacterial populations, occasionally driving them to local extinction and thereby releasing less competitive community members from competition (Alseth et al., 2024; Fazzino et al., 2020; Wei et al., 2011). Phage-mediated lysis also releases intracellular metabolites that can support the growth of non-host organisms and profoundly impact established metabolic interactions within microbial communities (Pherribo and Taga, 2023). However, phage-driven perturbations do not necessarily lead to bacterial host extinction. The emergence of genetic or phenotypic resistance, including mucoidy, together with spatial refuges and variation in host physiological state, can promote long-term coexistence between phages and susceptible bacteria (Chaudhry et al., 2020; Lourenço et al., 2020; Pyenson et al., 2024). Phage effects are therefore expected to be strongly dependent on community composition, resource availability, spatial structure, and the physiological state of bacterial hosts. In this context, the active replication detected during vegetable fermentation may represent not only a response to changes in host abundance, but also a process capable of redirecting microbial succession through selective mortality, competitive release, nutrient redistribution, and the selection of resistant subpopulations. Explicitly incorporating phage replication into models of fermented-food microbiomes may consequently improve our understanding of the variability, resilience and functional trajectories of these ecosystems.

## 6. Acknowledgements

This work was funded by the French National Research Agency (ANR) as part of the MetaSimFood project (ANR-21-CE21-0003). JCG received a PhD grant from the French National Research Agency (ANR) as part of the same project. We are grateful to the INRAE MIGALE bioinformatics facility (doi: 10.15454/1.5572390655343293E12) for providing computing and storage resources. We thank Lysiane Omhover Fougy, Marie-Helene Desmonts and Dalal Werner (Aerial, Illkirch, France) for providing the grated and sterilized vegetables.

## 7. Author contributions

**Julia Cantuti Gendre:** Conceptualization, Methodology, Formal analysis, Investigation, Writing - Original Draft. **Margareth Renault:** Methodology, Formal analysis, Writing – Review & Editing. **Anne-Cécile Leforestier:** Methodology, Formal analysis, Writing - Review & Editing. **Gwendoline Marquet:** Methodology, Writing - Review & Editing. **Françoise Irlinger:** Methodology, Writing - Review & Editing. **Nathan Fouchard:** Formal analysis, Writing - Review & Editing. **Sophie Landaud:** Validation, Writing - Review & Editing, Supervision. **Claire Le-Marrec:** Methodology, Writing - Review & Editing, Supervision. **Stéphane Chaillou:** Writing - Review & Editing, Project administration, Funding acquisition. **Eric Dugat-Bony:** Conceptualization, Writing - Review & Editing, Supervision, Project administration, Funding acquisition.

## 8. Data Availability Statement

Raw sequencing data for fermented vegetables metagenomes and viromes were deposited at the Sequence Read Archive (SRA) of the NCBI (https://www.ncbi.nlm.nih.gov/sra) as part of BioProject <u>PRJNA1230211</u>. The complete genome of L. mesenteroides CIM 3004 was deposited on Genbank under the accession number JCEMDN000000000. Short-reads from the genomic material of the insensitive mutant L. mesenteroides CIM 3004 BIM were deposited at the Sequence Read Archive (SRA) of the NCBI (https://www.ncbi.nlm.nih.gov/sra) as part of BioProject PRJNA1230211. All datasets are available through our National Research Data Platform (https://entrepot.recherche.data.gouv.fr/). Regarding the spontaneous fermentation experiment: microbial enumeration and pH data (https://doi.org/10.57745/DN4I5E); epifluorescence microscopy data (https://doi.org/10.57745/IPEUWS); shotgun metagenomics processed data (https://doi.org/10.57745/CEVLVN); viral metagenomics processed data (https://doi.org/10.57745/3EHB1F). Regarding the synthetic ecology experiment: (https://doi.org/10.57745/CBUOLW). All the scripts used for data analysis are available through Github (https://github.com/jcantuti/Phage_dynamic_fermented_food) and Zenodo (https://doi.org/10.5281/zenodo.20813383).

---

**Supplementary Figure S1: Experimental setup for experimental fermentation of cabbage and carrot in the laboratory**. 1) Grated vegetables (∼3.2 kg) and sterile sea salt (1.5% w/w) were placed in a sterile beaker under a biological safety cabinet. 2) The grated vegetables and sterile sea salt were thoroughly mixed in the sterile beaker under a biological safety cabinet, homogenized by hand using sterile gloves, and left to stand at room temperature for 1 hour. 3) The vegetable mixture was then separated from the exudate by pressing manually using sterile gloves and sterile gauze. 4 to 6) The vegetable and exudate were evenly distributed among individual jars, resulting in a final product weight of 135 g per jar. 7) A sterile plastic grid and a plastic pusher were placed in each jar to keep the vegetables submerged once the lid was sealed. 8) The sealed jars were then incubated at 20°C for fermentation.

**Supplementary Figure S2: Dynamic of the microbial community during spontaneous vegetable fermentation.** Relative abundance of reads per sample (%) at the domain level for cabbage (A) and carrot (B) samples.

**Supplementary Figure S3: Structural variations of the bacterial communities of fermented cabbage (A) and carrot (B).** Principal coordinate analysis (PCoA) was applied to the Bray-Curtis dissimilarity matrix derived from bacterial community profiles. Samples are colored according to the sampling day.

**Supplementary Figure S4: Structural variations of the viral communities of fermented cabbage (A) and carrot (B).** Principal coordinate analysis (PCoA) was applied to the Bray-Curtis dissimilarity matrix derived from viral community profiles. Samples are colored according to the sampling day.

**Supplementary Figure S5: Kinetics of acidification and changes in the microbial and viral load during carrot fermentation.** A) Evolution of lactic acid production during carrot fermentation. Concentrations are expressed in g/L of exudate. B) & C) Viable counts of Rahnella aquatilis CIRMBIA 2785 (B), Lactiplantibacillus plantarum CIRM-BIA 2443 (C). Bar indicates the average value of four replicates (independent laboratory-scale productions) in log CFU/g of vegetables, with error bars representing the standard deviation. The dashed red line indicates the detection limit for microbial enumeration (2 CFU/g for day 0 and 5 CFU/g for day 3 and 28). D) Concentration of Leuconostoc phage PLE001 in log PFU/g during carrot fermentation. The dotted red bar represents the limit of detection for phage titration set at 3.1 log CFU/g.

