## Supplementary figures for "Evidence of active phage replication during vegetable fermentation and implications for the dynamics of bacterial communities"

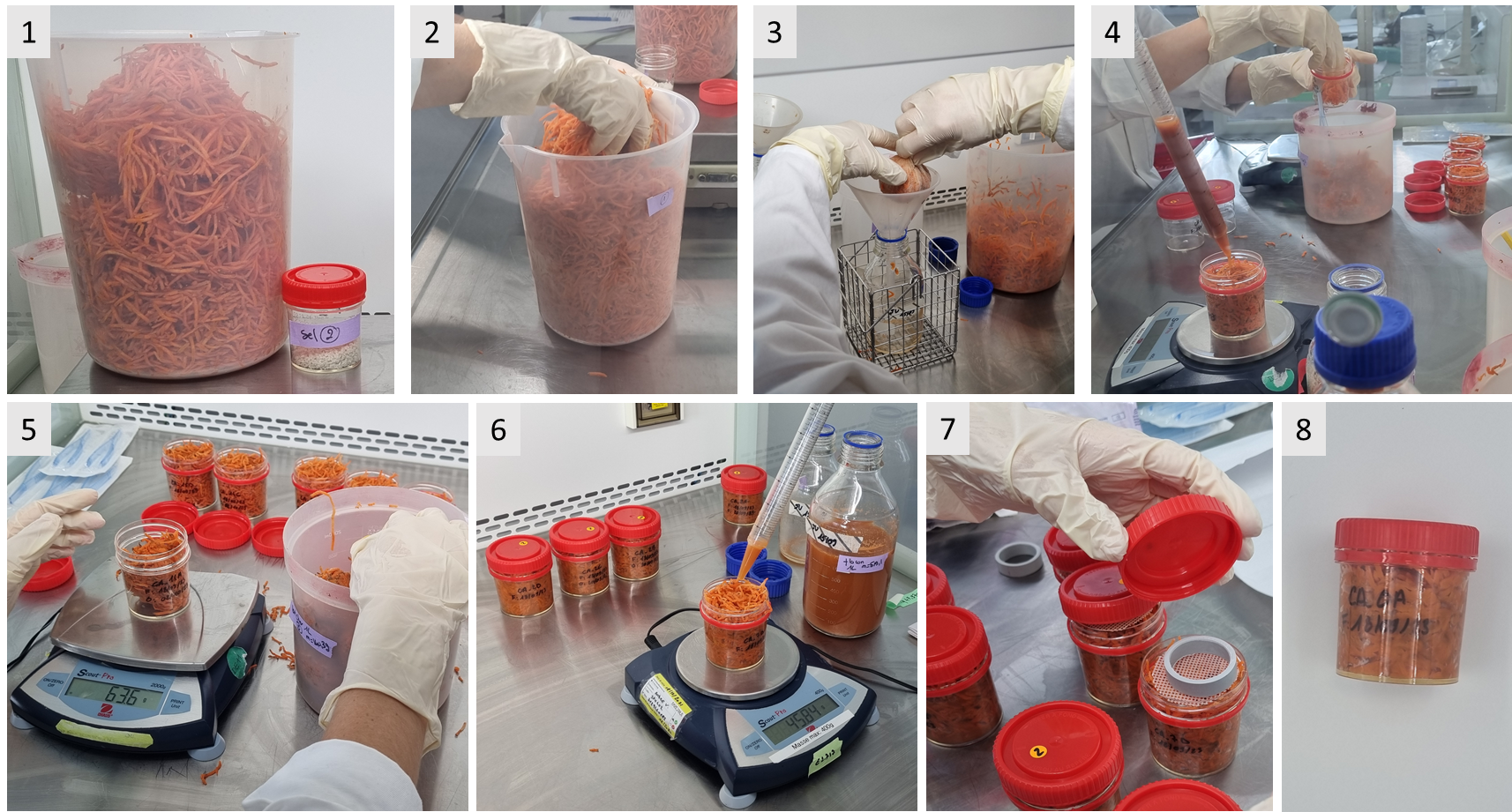

**Supplementary Figure S1: Experimental setup for experimental fermentation of cabbage and carrot in the laboratory.** 1) Grated vegetables (~3.2 kg) and sterile sea salt (1.5% w/w) were placed in a sterile beaker under a biological safety cabinet. 2) The grated vegetables and sterile sea salt were thoroughly mixed in the sterile beaker under a biological safety cabinet, homogenized by hand using sterile gloves, and left to stand at room temperature for 1 hour. 3) The vegetable mixture was then separated from the exudate by pressing manually using sterile gloves and sterile gauze. 4 to 6) The vegetable and exudate were evenly distributed among individual jars, resulting in a final product weight of 135 g per jar. 7) A sterile plastic grid and a plastic pusher were placed in each jar to keep the vegetables submerged once the lid was sealed. 8) The sealed jars were then incubated at 20°C for fermentation.

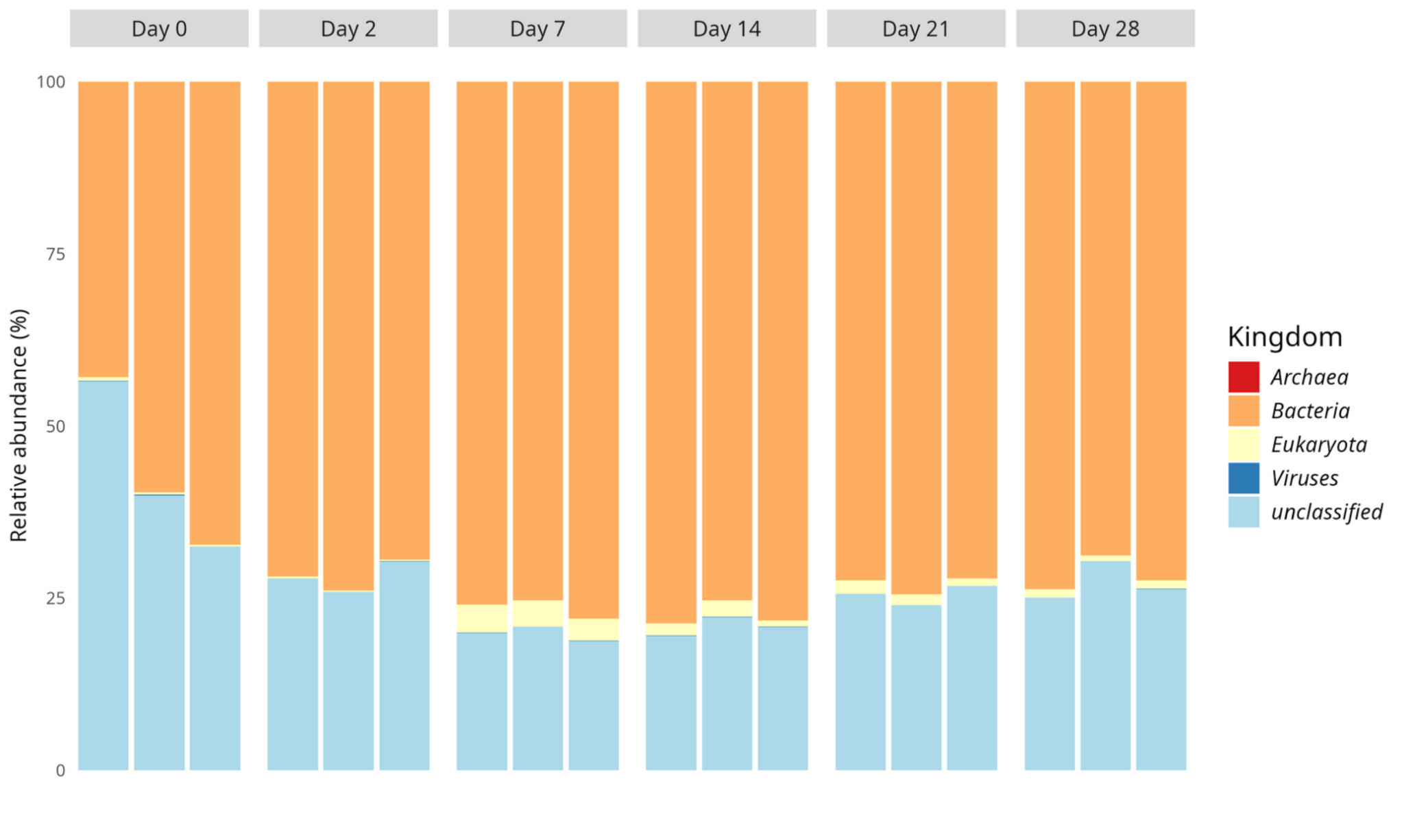

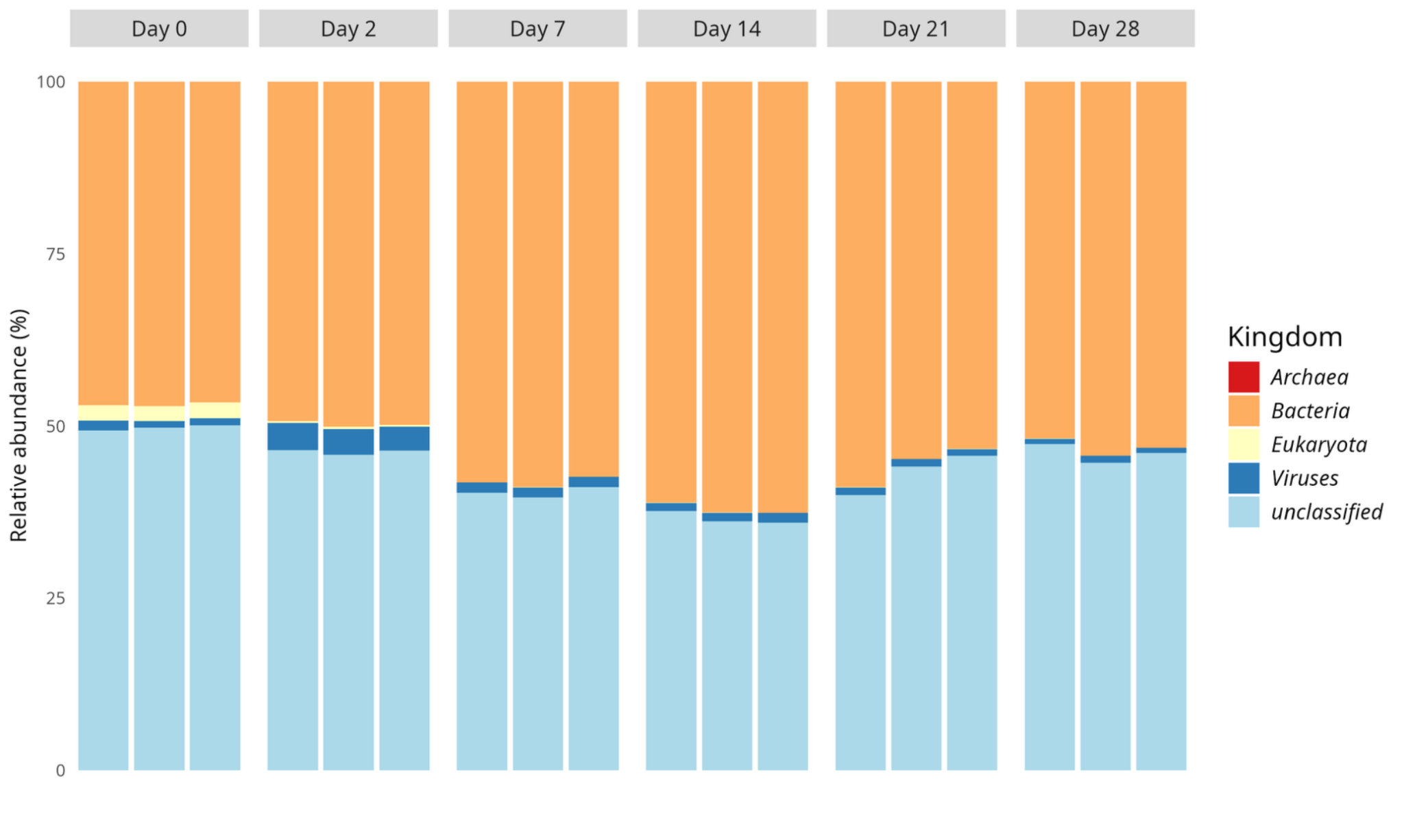

**Supplementary Figure S2: Dynamics of the microbial community during spontaneous vegetable fermentation.** Relative abundance of reads per sample (%) at the domain level for cabbage (A) and carrot (B) samples.

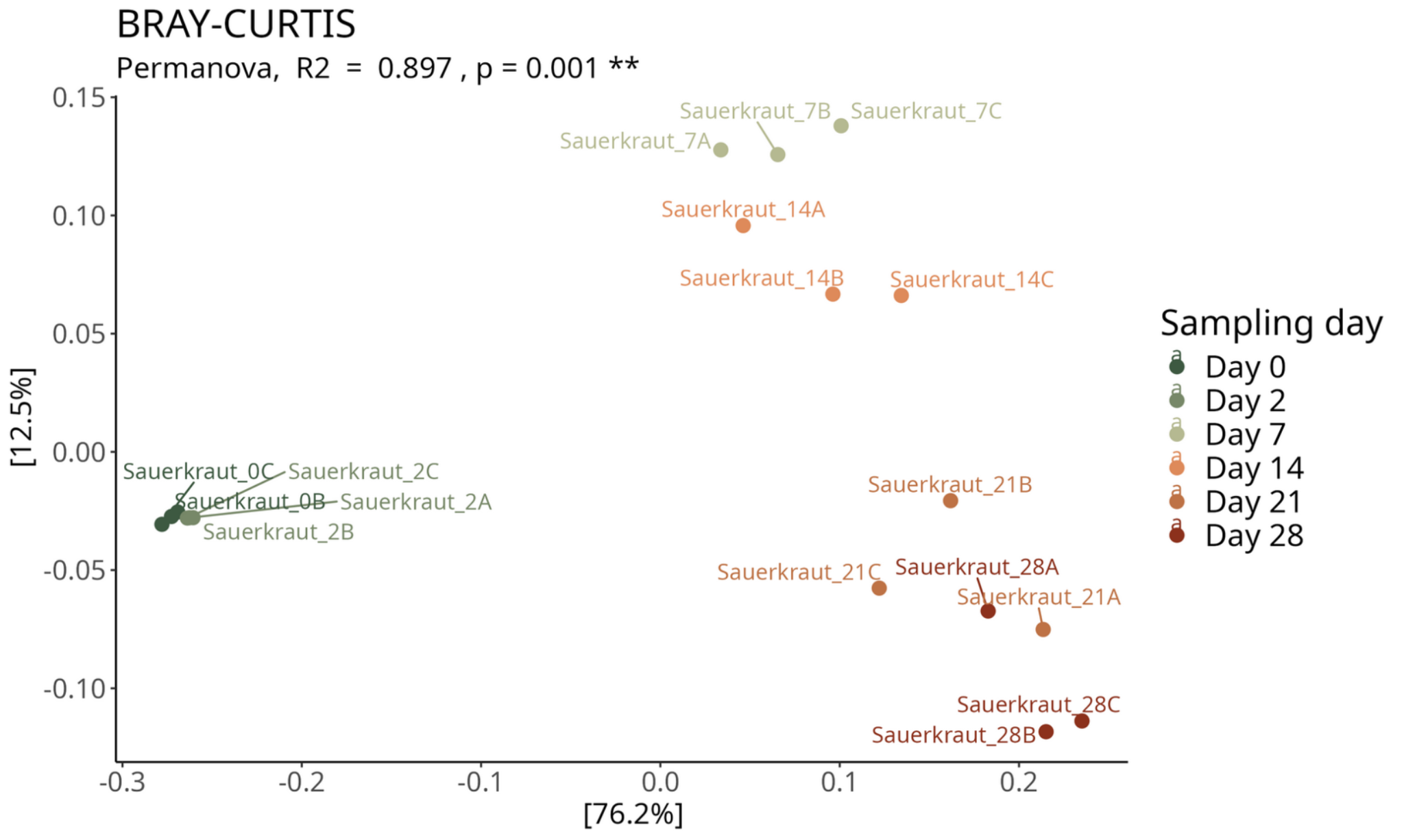

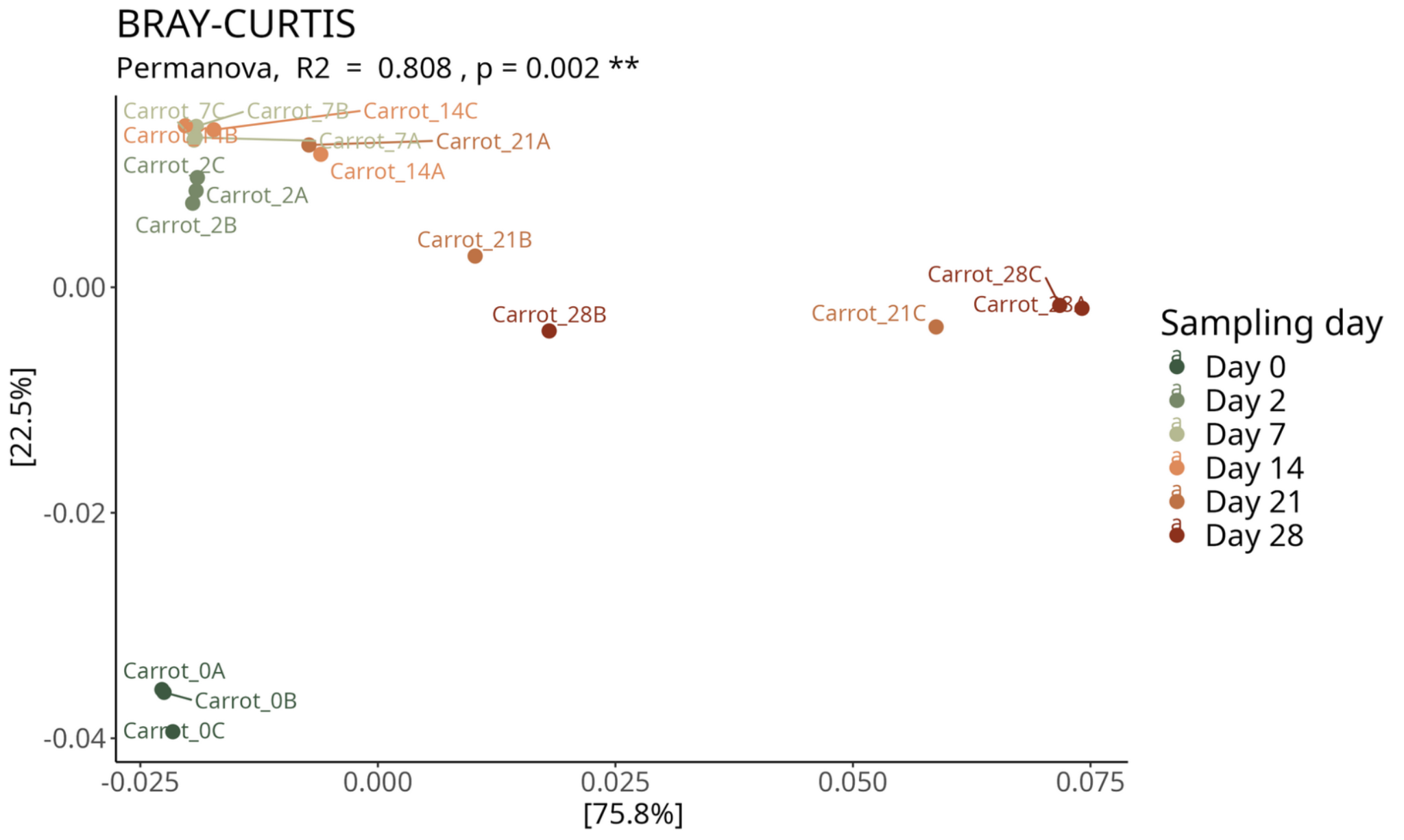

**Supplementary Figure S3: Structural variations of the bacterial communities of fermented cabbage (A) and carrot (B).** Principal coordinate analysis (PCoA) was applied to the Bray-Curtis dissimilarity matrix derived from bacterial community profiles. Samples are colored according to the sampling day.

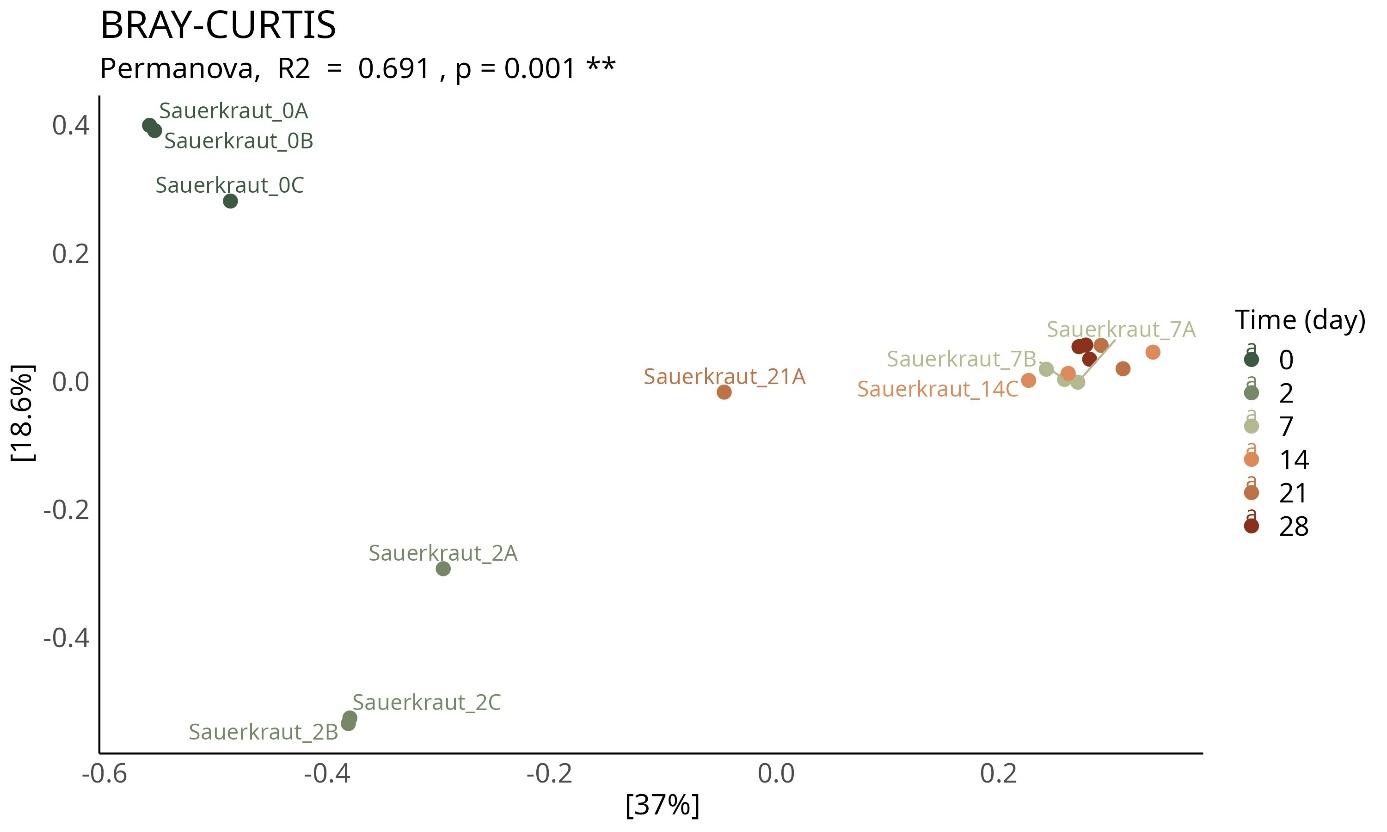

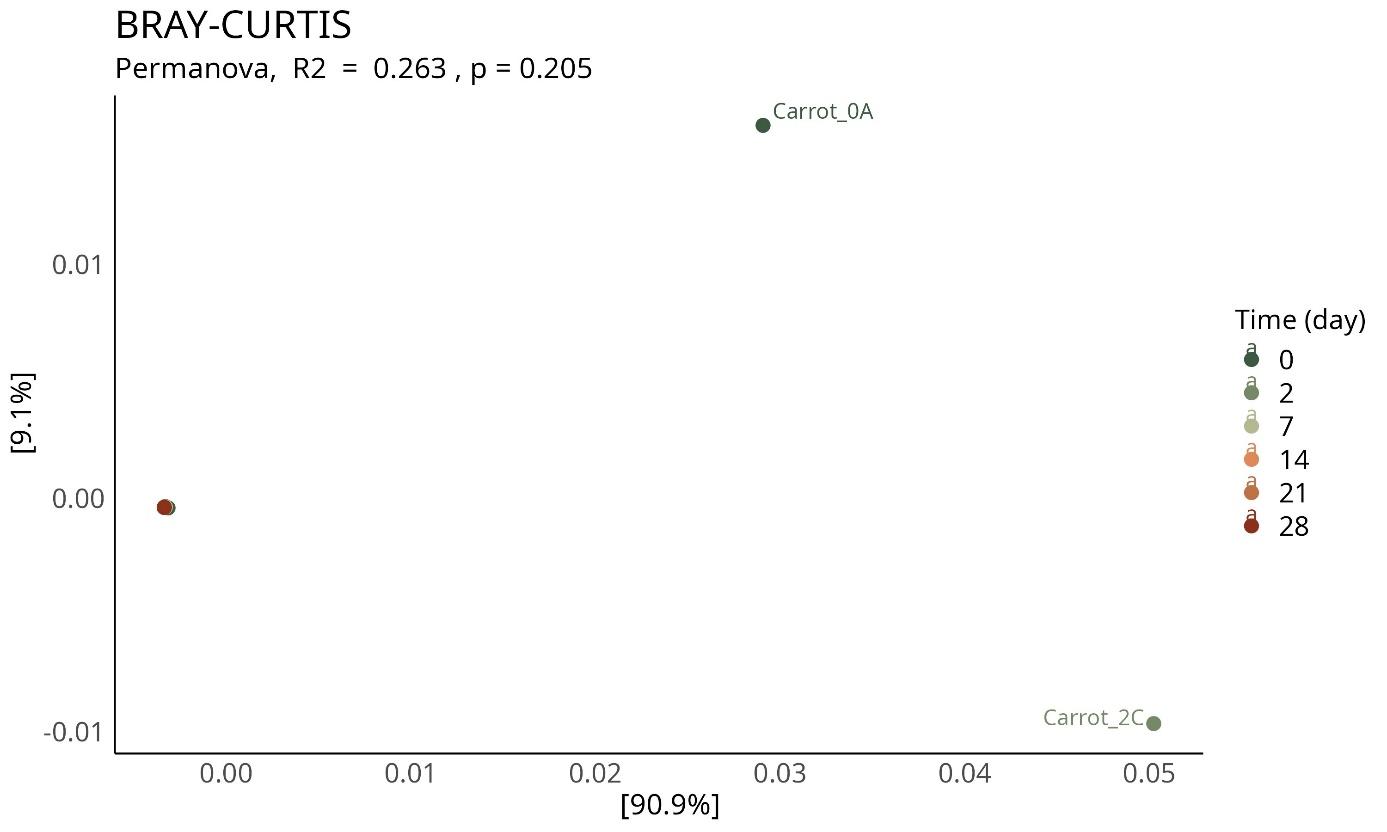

**Supplementary Figure S4: Structural variations of the viral communities of fermented cabbage (A) and carrot (B).** Principal coordinate analysis (PCoA) was applied to the Bray-Curtis dissimilarity matrix derived from viral community profiles. Samples are colored according to the sampling day.

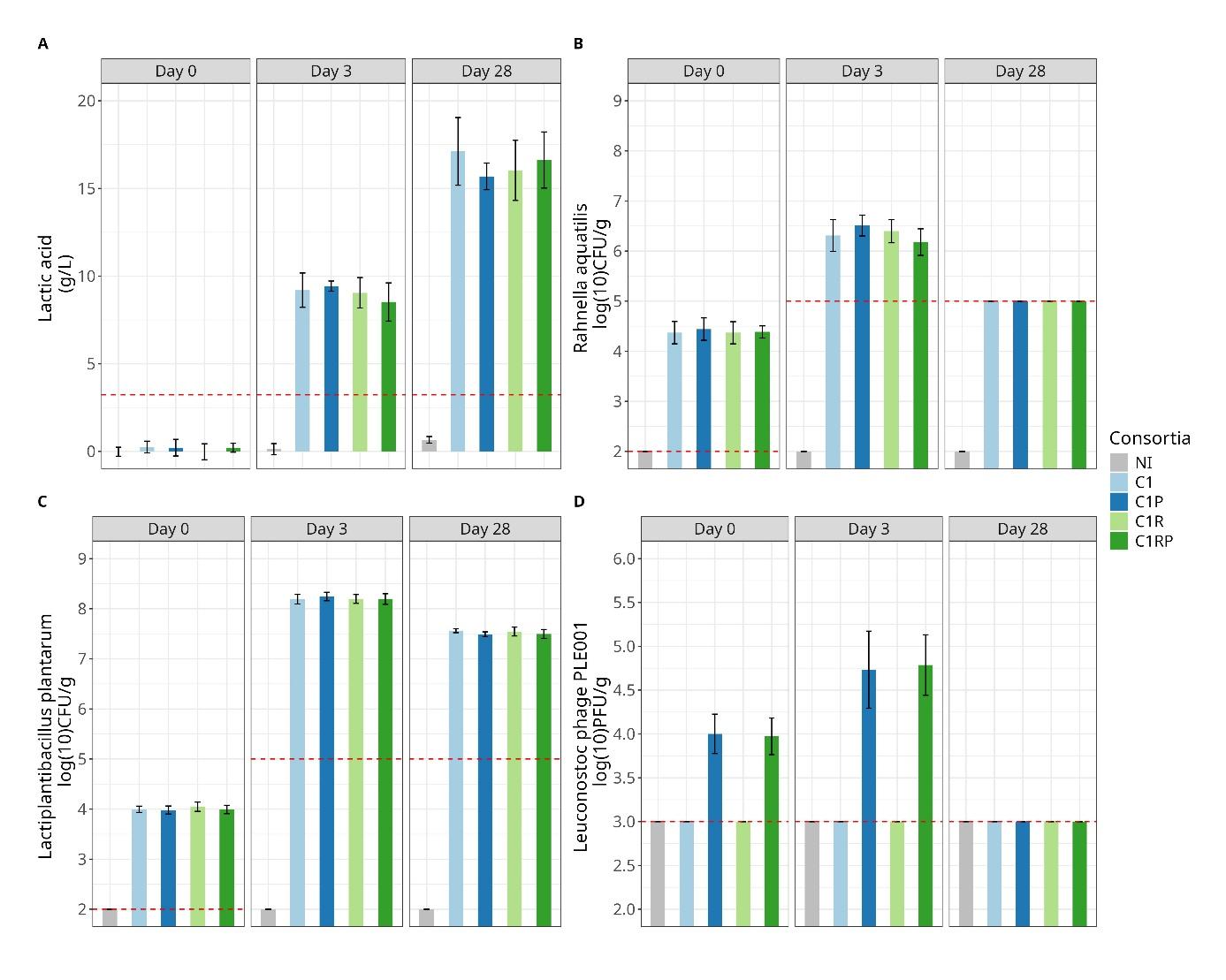

**Supplementary Figure S5: Kinetics of acidification and changes in the microbial and viral load during carrot fermentation**. A) Evolution of lactic acid production during carrot fermentation. Concentrations are expressed in g/L of exudate. B) & C) Viable counts of *Rahnella aquatilis* CIRMBIA 2785 (B), *Lactiplantibacillus plantarum* CIRM-BIA 2443 (C). Bar indicates the average value of four replicates (independent laboratory-scale productions) in log CFU/g of vegetables, with error bars representing the standard deviation. The dashed red line indicates the detection limit for microbial enumeration (2 CFU/g for day 0 and 5 CFU/g for day 3 and 28). D) Concentration of *Leuconostoc* phage PLE001 in log PFU/g during carrot fermentation. The dotted red bar represents the limit of detection for phage titration set at 3.1 log CFU/g.
